# Hypoxia-inducible-factors differentially contribute to clinical disease and viral replication during RSV infection

**DOI:** 10.1101/2023.08.15.553422

**Authors:** Dorothea R. Morris, Yue Qu, Aline Haas de Mello, Yava L. Jones-Hall, Tianshuang Liu, Meredith Weglarz, Teodora Ivanciuc, Roberto P. Garofalo, Antonella Casola

**Affiliations:** Department of Microbiology & Immunology, The University of Texas Medical Branch, Galveston, TX, USA; School of Population & Public Health, The University of Texas Medical Branch, Galveston, TX, USA; Department of Pediatrics, The University of Texas Medical Branch, Galveston, TX, USA; School of Veterinary Medicine and Biomedical Science, Texas A&M University, College Station, TX, USA

**Author notes:** Correspondence: Antonella Casola.

**Keywords:** RSV, respiratory syncytial virus, hypoxia-inducible-factors, HIF-1a, HIF-2a, PX478, PT2385

## Abstract

Hypoxia-inducible-factors (HIF) are transcription factors that regulate cellular adaptation to hypoxic conditions, enabling cells to survive in low-oxygen environments. Viruses have evolved to activate this pathway to promote successful viral infection, therefore modulation of HIFs could represent a novel antiviral strategy. In previous *in vitro* studies, we found that respiratory syncytial virus (RSV), a leading cause of respiratory illness, stabilizes HIFs under normoxic conditions, with inhibition of HIF-1α resulting in reduced viral replication. Despite several HIF modulating compounds being tested/approved for use in other non-infectious models, little is known about their efficacy against respiratory viruses using relevant animal models. This study aimed to characterize the disease modulating properties and antiviral potential of HIF-1α (PX478) and HIF-2α (PT2385) inhibitors in RSV-infected BALB/c mice. We found that inhibition of HIF-1α worsen clinical disease parameters, while simultaneously improving lung inflammation and airway function. Additionally, blocking HIF-1α resulted in significantly reduced viral titer at early and peak time points of RSV replication. In contrast, inhibition of HIF-2α was associated with improved clinical parameters, with no changes in airway function, enhanced immune responses and reduced early and peak lung viral replication. Analysis of lung cells found significant modification in the T-cell compartment that correlated with changes in lung pathology and viral titers in response to each HIF inhibitor administration. This study underscores the differential roles of HIF proteins in RSV infection and highlights the need for further characterization of the compounds that are currently in use or under therapeutic consideration.

## 2 Introduction

Respiratory syncytial virus (RSV) is a prominent cause of respiratory infections among infants, young children, and the elderly. Each year, there are an estimated 33 million infections and 150,000 deaths [1]. Infection with RSV often results in damage to the airways and a dysregulated immune response that contributes to the development of bronchiolitis and pneumonia. Although prophylactic treatment and vaccines licensed for use in pregnant women and elderly patients are available, there are no therapeutic agents licensed for the treatment of RSV infection [2–4]. Furthering our understanding of the pathology of RSV is important to protecting these vulnerable populations.

As an obligate intracellular pathogen, viruses rely on host-cell machinery to establish productive infections. Many viruses have been shown to modify cellular metabolism to achieve virion assembly by increasing sources of energy and generating pools of free nucleotides, amino acids, lipid materials, and glycoproteins [5]. One mechanism frequently associated with this is the manipulation of the hypoxia-inducible-factor (HIF) pathway. As the gatekeeper to a diverse biological network, the use of the HIF pathway allows many viruses to create a cellular environment that favors viral replication, persistence, and immune evasion. The manipulation of this pathway has been observed with several DNA and RNA viruses, even under oxygen-rich conditions [6, 7]. The significance of HIF in cancer pathology has led much of the existing literature to focus on viruses known to induce tumor formation as part of their pathogenic mechanism [6, 7]. Surprisingly, limited research has explored the clinical manifestations and pathological contributions of HIFs to viral respiratory infections [6–9].

HIFs are transcription factors that regulate the expression of a wide range of target genes in response to low oxygen levels [10]. These target genes have crucial roles in diverse biological processes, enabling cells to adapt and survive under hypoxic conditions. The HIF complex is a heterodimer consisting of an inducible α subunit (HIF-1α, HIF-2α, or HIF-3α) and a constitutively expressed β subunit (HIF-1β) [11]. While there is overlap in their functionality, the HIF-α isoforms also exhibit distinct transcriptional regulation in response to hypoxia and disease states [12–14]. HIF-1α is the most extensively studied isoform of the HIF family. Expressed in nearly all cell types, HIF-1α is the master regulator of cellular and systemic responses to hypoxia. Genes activated by HIF-1α modify a broad range of processes including energy metabolism, angiogenesis, epithelial-mesenchymal transition (EMT), and immune cell regulation [12, 14]. HIF-2α, though not as thoroughly described as HIF-1, has been shown to exhibit a more limited distribution; predominately expressed in specific organs such as the kidney, liver, and lung. Genes upregulated by HIF-2α in models of cancer and of non-infectious tissue repair are primarily involved in erythropoiesis, vascular development, and iron metabolism [14]. HIF-3α is the most recent of the HIF isoforms to be identified. Because of this, the precise distribution and function of HIF-3α is still being investigated.

Our laboratory recently found that RSV stabilizes both HIF-1α and HIF-2α in primary small airway epithelial cells and diverts core metabolic activity towards the glycolytic pathway in a HIF-1α dependent manner [15]. Inhibition of HIF-1α, but not HIF-2α, was associated with reduced viral replication. Given the multiple roles of HIFs in both the epithelial and immune compartments, we investigated their involvement in RSV lung disease and pathology by utilizing established inhibitors of HIF-1α (PX478) and HIF-2α (PT2385) in a BALB/c mouse model. Our results underscore the distinct functionality of the individual HIF-α subunits in the development of RSV pathogenesis and indicate the importance of testing HIF therapeutics approved for conditions such as cancer in complex viral respiratory infection biological models for potential benefits, but also unwanted side effects.

## 3 Materials and methods

### 3.1 Animal experiments

Female, 8 to 10-week-old BALB/c mice were purchased from Envigo (Indianapolis, IN, USA). Mice received either PX478 or PT2385 (HY-10231 and HY-12867, MedChemExpress, Monmouth Junction, NJ, USA) by oral gavage at a dose of 20 or 50 mg/kg, respectively. PX478 was dissolved in PBS, while PT2385 was dissolved in 10% DMSO + 90% Corn Oil (CO). For anti-HIF1α, PX478 treated mice were separated into three groups. The first received PBS as the control group. The second received PX478 every 24 h for 10 days (PXQD). The third group received PX478 every 48 h for 10 days (PXQAD). For anti-HIF2α, mice received either CO or PT2385 every 24 h for 10 days. Within 15 minutes of the first treatment, mice were anesthetized and intranasally (IN) inoculated with 50μL of either PBS or RSV Long Strain at a dose of 5×10^6^ plaque forming units (PFU). Preparation of RSV Long Strain (ATCC, Manassas, VA, USA) was done as described previously [16]. All mice were monitored over the 10-day infection period for changes in bodyweight loss and clinical illness score on a 0-to-5 grading scale (0 = healthy, 1 = barely ruffled fur, 2 = ruffled fur but active, 3 = ruffled fur and inactive, 4 = ruffled fur, inactive and hunched, 5 = dead). All care and procedures involving mice in this study were completed in accordance with the recommendations in the *Guide for the Care and Use of Laboratory Animals* of the National Institutes of Health and the UTMB institutional guidelines for animal care. The Institutional Animal Care and Use Committee (IACUC) of UTMB approved these animal studies under protocol 9001002.

### 3.2 RNA extraction and reverse transcription-quantitative PCR (RT-qPCR)

Lung tissue was homogenized in TRIzol reagent (15596018; Thermo Fisher Scientific, Waltham, MA, USA) and the RNA was isolated using a combination of TRIzol-based method and the RNeasy Mini Kit (74104; Qiagen, Hilden, Germany). Briefly, after phase separation with chloroform, the top aqueous layer was further processed using the Qiagen RNeasy Mini kit spin columns by following manufacturer’s protocol. On-column DNase digestion was performed with the RNase-free DNase set (79254; Qiagen). cDNA synthesis was performed with 1 µg of RNA using iScript Reverse Transcription Supermix (1708841; Bio-Rad Laboratories, Hercules, CA, USA). The cDNA was diluted four times with nuclease-free water, and the qPCR was done using 4 µL of cDNA, pre-mixed probe and primers (TaqMan Gene Expression Assays, Applied Biosystems, Waltham, MA, USA), and TaqMan Universal Master Mix (4440040; Applied Biosystems). The Custom Plus TaqMan RNA Assay (4441114, Applied Biosystems) was used to assess the expression of *RSV N* (Assay ID: ARU66XH). The following mouse TaqMan Gene Expression Assays (4331182, Applied Biosystems) were used: *Hif1α* (Mm00468869_m1), *Pgk1* (Mm00435617_m1), *Hk2* (Mm00443386_m1), *Slc2a1* (Mm00441480_m1), *Serpine1* (Mm00435858_m1), *Epo* (Mm01202755_m1) and *Epas1* (Mm01236112_m1). The Eukaryotic 18S rRNA Endogenous Control (4352930E, Applied Biosystems) was used for normalization. The qPCR assays were run in the Bio-Rad CFX Connect real-time system. The delta-delta Ct method was used to calculate the relative changes in gene expression.

### 3.3 Bronchoalveolar lavage and viral replication

The BAL fluid was obtained as previously described [17]. A small aliquot was used to determine the total cell count and cellular differentiation of the BAL by cytospin analysis. The remaining BAL fluid was centrifuged, and the supernatants were collected and stored at -80C. To assess viral replication, lungs were collected from mice at days 2, 4, 7, and 10 post-infection (p.i.) to perform plaque assays, as previously described [17].

### 3.4 Flow cytometry

At days 1, 2, 4 and 7 p.i., mice were euthanized and the whole lung was collected. The tissue was minced and digested with 10mg/mL DNase I (Sigma-Aldrich, St. Louis, MO, USA) and 50mg/mL collagenase IV (Worthington Biochemical, Lakewood, NJ, USA). The tissue was incubated at 37C for 30 minutes and then passed through a 70-μm cell strainer in RPMI 1640 medium + 10% FBS (CRPMI) and red blood cells were removed by using Red Blood Cell Lysis Buffer (Sigma-Aldrich, Burlington, MA, USA). TruStain FcX (Biolegend, San Diego, CA, USA) was first used to reduce non-specific binding, followed by Zombie NIR Viability dye. To assess cellular populations of neutrophils, alveolar macrophages, and natural killer (NK) cells at days 1 and 2 p.i. a comprehensive panel of fluorochrome-labeled antibodies was utilized. The antibodies employed included FITC-anti-CD45 (clone 30-F11), BV785-anti-CD11b (clone M1/70), PerCP/Cyanine 5.5-anti-Ly6G (clone 1A8), BUV395-anti-CD11c (clone HL3), PE-Cy7-anti-F4/80 (clone BM8), BV510-anti-MHCII (clone M5/114.15.2), BV711-anti-CD3ε(clone 145-2C11), and PE/Dazzle 594-anti-CD49b (clone DX5). For the analysis of CD4^+^ and CD8^+^ T-cell populations at days 4 and 7 p.i. the panel included: FITC-anti-CD45 (clone 30-F11), BV785-anti-CD3 (clone 17A2), AF 700-anti-CD4 (clone GK1.5), BV510-anti-CD8a (clone 53-6.7), PerCP/Cy5.5-anti-CD44 (clone IM7), BV605-anti-CD62L (clone MEL-14), and PE/Cy7-anti-CD69 (clone H1.2F3). All antibodies were purchased from Biolegend, Thermo Fisher Scientific, Novus Biologicals, or R&D Systems and titrated prior to use. Data were acquired on a BD FACSymphony A5 SE in the UTMB Flow Cytometry and Cell Sorting Core and analyzed using FlowJo software version 10.9.0 (BD Bioscience, Franklin Lakes, NJ, USA).

### 3.5 Assessment of airway function and lung histopathology

Bronchoconstriction was evaluated by measuring Penh values in unrestrained mice using whole-body barometric plethysmography (Buxco, Troy, NY, USA) as previously described [18]. Total protein was measured using BioRad Total Protein Dye (Bio-Rad Laboratories, Hercules, CA, USA). For histology, the left lung was collected from mice at days 4, 7, and 10. The tissue was fixed in 10% neutral buffered formalin and subjected to paraffin embedding. Sections were cut and stained with H&E and evaluated by a board-certified pathologist with expertise in mouse lung (Y.L.J.-H). Assessment of the tissue was done as previously described [19].

### 3.6 Statistical analysis

Statistical analyses were performed using a two-way mixed ANOVA or an unpaired student’s t-test as appropriate. Analyses were done using GraphPad Prism 9.5.1 (GraphPad Software, Inc., San Diego, CA, USA). Results are expressed as mean ± SEM and p ≤ 0.05 value was selected to indicate significance.

## 4 Results

### 4.1 Effect of PX478 administration on RSV infection associated disease parameters and viral replication

To investigating the role of HIF-1α in RSV disease, groups of BALB/c mice were initially administered PX478 at a dose of 20 mg/kg by oral gavage every 24 h (PXQD) over the 10-day infection period (Fig. S1A). Shortly after the first dose of PX478, mice were anesthetized and intranasally (IN) infected with RSV at a dose of 5×10^6^ PFU. We found that PXQD treatment was associated with worsening body weight and illness score, characterized by an inability to regain weight regardless of infection status (Fig. S1B, S1C).

To assess airway function during RSV infection, we measured bronchoconstriction using whole-body plethysmography, as previously described [17]. Uninfected mice that received PX478 showed no significant alterations of Penh values at any timepoint tested, compared to the PBS/PBS control group. RSV infected PX478-treated mice showed significant improvement in Penh at days 1, 5, and 6 p.i., with values comparable to the PBS/PBS control mice throughout the disease course (Fig. S1D).

We next measured total protein level in the BAL fluid as an indicator of epithelial damage. PX478 treatment did not alter BAL protein levels in uninfected mice, while RSV infected PX478-treated mice showed significant improvements in total protein at day 4 p.i., compared to the RSV/PBS control mice (Fig. S1E).

To assess the effect of PX478 treatment on viral replication, we performed plaque assays at early and peak timepoints of day 2 and 4 p.i.. We found that PX478 treatment significantly reduced replication of days 2 and 4 p.i., when compared to the RSV/PBS control mice. However, RSV replication in RSV PX478-treated mice persisted through day 7 p.i., a timepoint when RSV/PBS control mice had achieved viral clearance, with some mice having detectable replication in the lung tissue as late as day 10 p.i. (Fig. S1F).

To improve the toxicity observed on body weight loss and illness in the PBS/PXQD treated mice, we tested lower doses of the compound (5 to 15 mg/kg) for changes in body weight and viral replication. Following the same treatment schedule, we found bodyweight to be improved for some of the doses in PBS mice treated with PX478, but none of the RSV mice treated with PX478 had comparable antiviral activity (data not shown). The administration of PX478 at 20 mg/kg was then changed to every 48 h (PXQAD) over the 10-day infection period (Fig. 1A). Uninfected mice that received the compound on alternate days no longer exhibited substantial weight-loss or heightened illness in comparison to the PBS/PBS control mice. However, RSV infected mice treated with PXQAD continued to demonstrate significantly worse body weight-loss and illness score in comparison to the RSV/PBS mice, although these mice showed signs of recovery at the later stages of the disease course (Fig. 1B, 1C).

**Figure 1.**
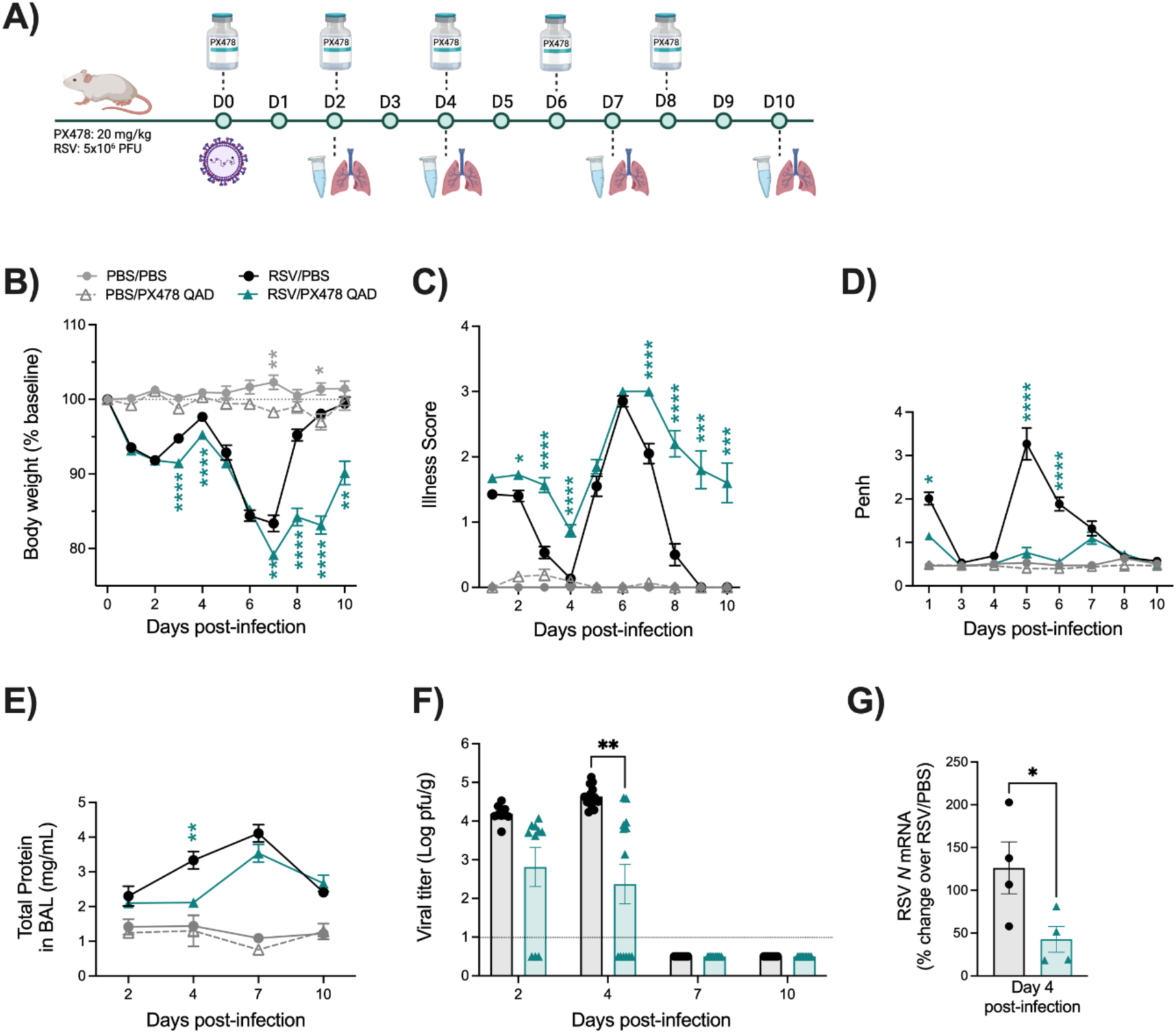
Assessment of clinical disease, airway function, and viral replication following HIF-1α inhibition during RSV infection. The experimental design for mice treated with PX478 QAD is described in (A). Following treatment, all mice were monitored daily for changes in (B) body weight and (C) illness score. (D) Bronchoconstriction, represented by baseline Penh, was measured by plethysmography. At days 2, 4, 7, and 10 p.i., (E) total protein was measured in the BAL fluid and the lung was collected for assessment of (F) viral replication by plaque assay. At day 4 p.i, (G) RSV *N* mRNA levels were measured by RT-qPCR. For clinical disease, data are pooled from four independent experiments (PBS/PBS n = 6-8, PBS/PX n = 6-18, RSV/PBS n = 10-30, RSV/PX n = 9-30 mice/group). For bronchoconstriction, data are pooled from three independent experiments (PBS/PBS and PBS/PX n = 6-12, RSV/PBS and RSV/PX n = 6-30 mice/group). For total protein, data are pooled from two independent experiments (PBS/PBS and PBS/PX n = 4-6, RSV/PBS and RSV/PX n = 8-10 mice/group). For viral replication by plaque assay, data are pooled from two independent experiments (n = 10-14 mice/group). For RT-qPCR data are from one independent experiment (n = 4 mice/group). Data are expressed as mean ± SEM. Significant results were determined by two-way mixed ANOVA (B-F) and unpaired t test (G) (* p ≤ 0.05, ** p ≤ 0.01, *** p ≤ 0.001, **** p ≤ 0.0001).

Lung function was assessed by measuring bronchoconstriction (Penh) and BAL total protein levels. RSV infected mice that received PXQAD had significant improvements in Penh at days 1, 5, and 6 p.i., with values comparable to the PBS/PBS control mice throughout the disease course (Fig. 1D). RSV/PXQAD mice also had significant improvement in BAL total protein level at day 4 p.i. as compared to the RSV/PBS control mice (Fig. 1E).

To assess the effect of PXQAD treatment on viral replication, we performed plaque assays at early and peak timepoints of day 2 and 4 p.i, as well as days 7 and 10 p.i. to assess viral clearance. RSV/PXQAD mice showed significantly reduced RSV replication at day 4 p.i., with no significant difference in viral clearance at days 7 and 10 p.i. when compared to the RSV/PBS group (Fig. 1F). The change in peak RSV replication was further supported by reductions in the RSV *N* gene by RT-qPCR at day 4 p.i. (Fig. 1G). All subsequent experiment described below were then performed using the PXQAD treatment protocol.

### 4.2 Effect of PX478 administration on RSV-induced HIF-1α target genes

To confirm inhibition of RSV-induced HIF-1α activation by PX478 treatment, we analyze the expression of *Hif1α* and the HIF-1α-target genes phosphoglycerate kinase 1 (*Pgk1*), hexokinase 2 (*Hk2*), and solute carrier family 2 (facilitated glucose transporter) member 1 (*Slc2a1*), at day 2 p.i. in lungs of mice untreated or treated with the compound (Fig. 2). RSV infection resulted in upregulation of HIF-1α and HIF-1α-target genes, which was significantly reduced by PX478 treatment. Specificity of PX478 was determined by assessment of the HIF-2α−dependent genes erythropoietin (*Epo*) and *Serpine1*. *Epo* was not detectable in the lung tissue of any mice. RSV infection induced the expression of *Serpine1*, which was not changed by PX478 treatment, indicating specificity of the compound.

**Figure 2.**
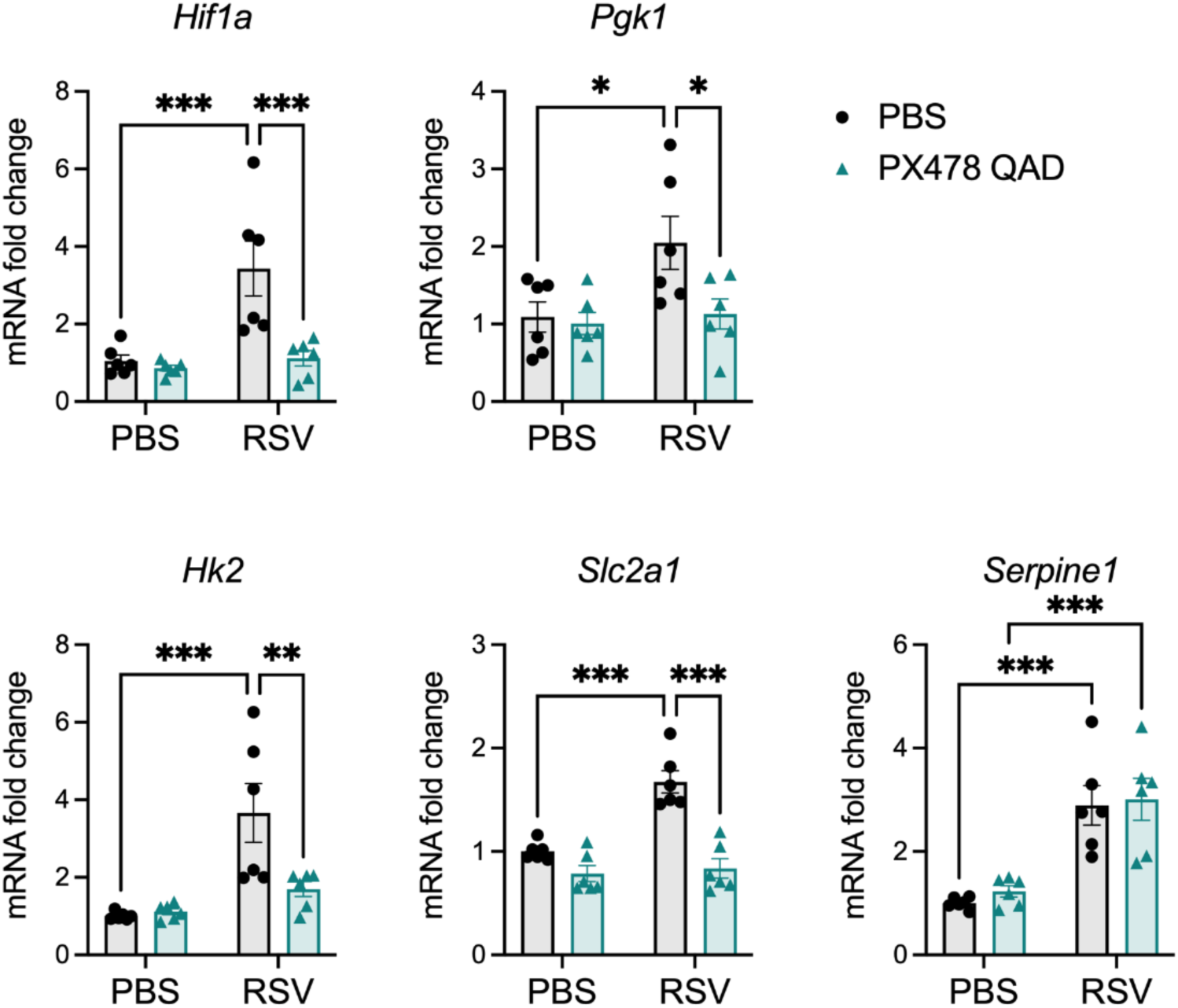
Effect of PX478 on Hif1a and HIF-1α and -2α target genes in the lungs of mice during RSV infection. Mice were euthanized at day 2 p.i. and RNA was extracted from the lung tissue. *Hif1a*, the HIF-1α targets *Pgk1*, *Hk2*, and *Slc2a1* (GLUT1), and the HIF-2α target *Serpine1* (PAI-1) gene expression was measured by RT-qPCR (n = 6 mice/group). Data are expressed as mean ± SEM. Significant results were determined by two-way ANOVA followed by Bonferroni’s multiple comparisons test (* p ≤ 0.05, ** p ≤ 0.01, *** p ≤ 0.001).

### 4.3 Inhibition of HIF-1α leads to reduced immune responses following RSV infection

The immune response to an RSV infection plays a key role in determining disease severity and the control of RSV replication [20]. To understand the impact of HIF-1α inhibition on the immune response to RSV, we first evaluated the cellular composition of the BAL fluid. Uninfected mice that received PXQAD showed no significant changes of BAL cellular composition as compared to the PBS/PBS control mice (Fig. 3A-D). On the other hand, RSV infected mice that received the compound demonstrated significant reductions in the number of total cells present in the BAL (Fig. 3A), specifically in the number of lymphocytes (Fig. 3B) and macrophages (Fig. 3C) at all time points tested, compared to RSV/PBS mice. Neutrophil cell counts were also significantly reduced at day 2 p.i. but remained similar to the RSV/PBS control mice at all other time points (Fig. 3D). Together, these data suggest HIF-1α is important for the recruitment of immune cells within the airway space.

**Figure 3.**
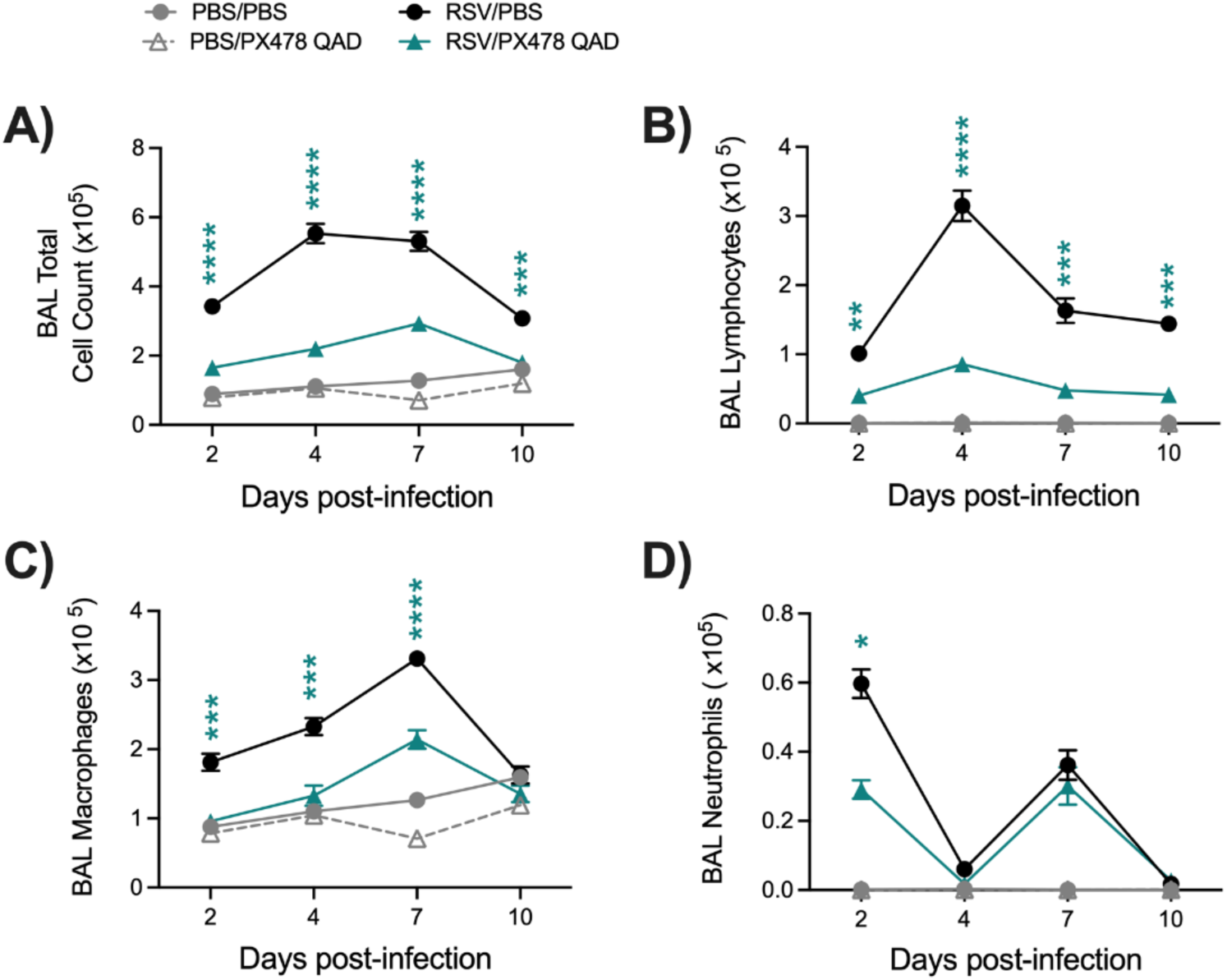
Differential cell count of the BAL fluid following HIF-1α inhibition during RSV infection. At days 2, 4, 7, or 10 p.i., BAL fluid was collected from mice that had been treated or not with PX478 QAD. This BAL was used to obtain (A) total cell counts, as well as differential cell counts consisting of (B) lymphocytes, (C) macrophages, and (D) neutrophils. Data are pooled from two independent experiments (PBS/PBS and PBS/PX478 n = 4-6, RSV/PBS and RSV/PX478 n = 10 mice/group). Data are expressed as mean ± SEM. Significant results were determined by two-way mixed ANOVA (* p ≤ 0.05, ** p ≤ 0.01, *** p ≤ 0.001, **** p ≤ 0.0001).

CD4^+^ and CD8^+^ T-cells play a fundamental role in controlling RSV replication and clearance within the host [20–22]. With reduced lymphocyte numbers in the BAL and the complex outcome to viral replication (prolonged viral replication in the PXQD treatment schedule), we next wanted to assess the impact of HIF-1α inhibition on the T-cell population within the lung tissue. To do so, flow cytometry analysis was performed using whole lung tissue at days 4 and 7 p.i. as depicted in Fig. S2. RSV infected mice treated with PXQAD demonstrated a significant reduction of total leukocytes (Fig. 4A), CD4^+^ (Fig. 4B) and CD8^+^ (Fig. 4C) T-cells present in the lungs at days 4 and 7 p.i., compared to the RSV/PBS control mice. We then further examined CD4^+^ T-cell subpopulations including naïve, effector, and active cells. We found the absolute cell numbers for all three CD4^+^ subpopulations to be significantly reduced at day 4 p.i. (Fig. 4D and Table S1). In accordance, the percentage of effector and active CD4^+^ cells were decreased while the percent of naïve CD4^+^ cells increased as compared to the RSV/PBS control mice (Table S1). At day 7 p.i., the absolute cell counts of effector CD4^+^ T-cells remained significantly reduced in the lung tissue of RSV/PXQAD mice. The absolute cell counts for naïve and active CD4^+^ T-cells as well as the percent change for all three CD4^+^ subpopulations were comparable to the RSV/PBS control mice at day 7 p.i. (Fig. 4D and Table S1).

**Figure 4.**
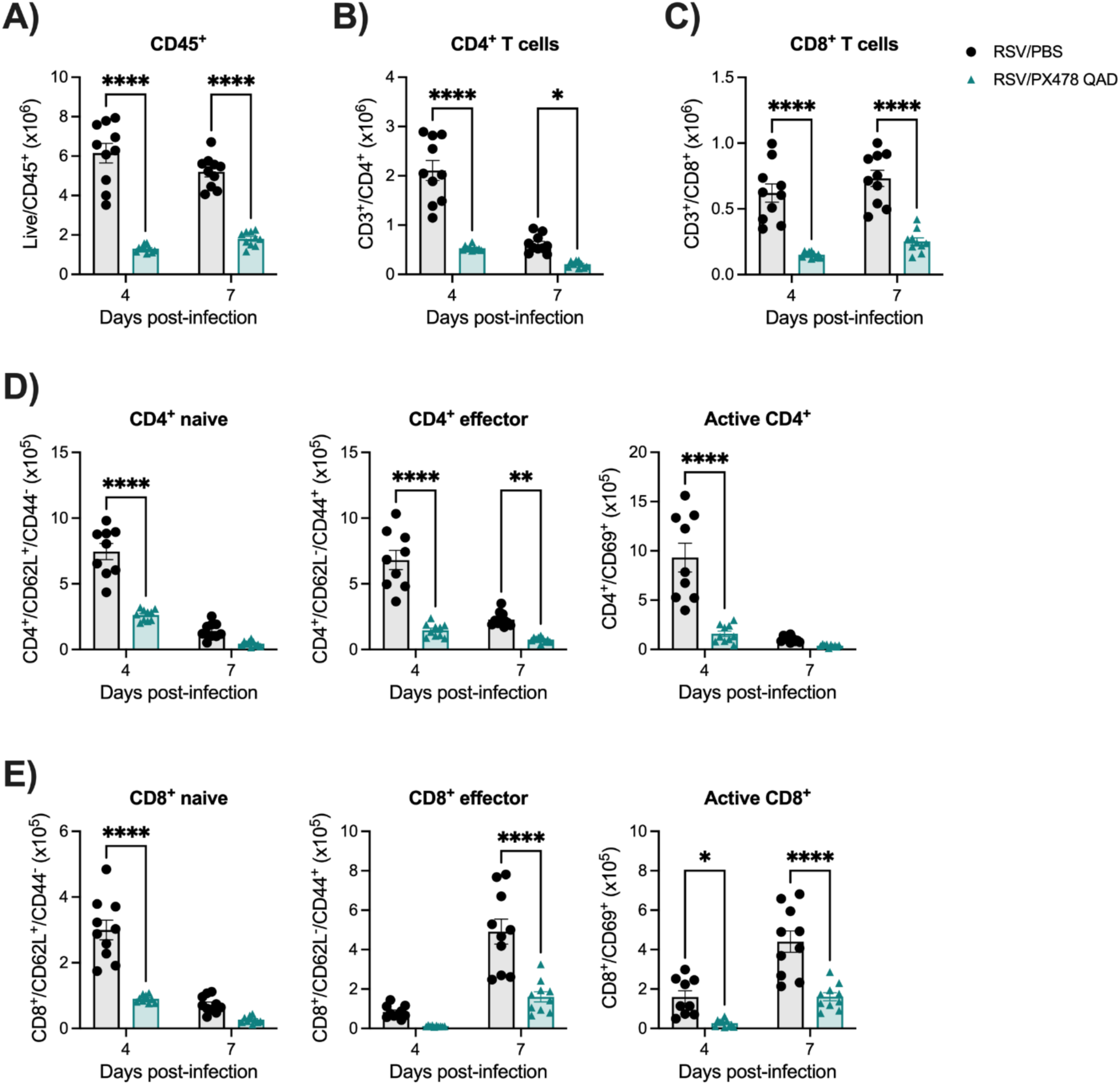
Assessment of CD4^+^ and CD8^+^ T cells following HIF-1α inhibition during RSV infection. Whole lung tissue was collected at peak RSV replication and a timepoint of viral clearance, days 4 and 7 p.i., respectively. Single cell suspension was prepared, stained with live/dead cell dye and fluorochrome-conjugated antibodies, and acquired by flow cytometry. Data were analyzed following the gating strategy in Figure S2. Quantification of absolute cell counts of (A) Live CD45^+^, (B) CD4^+^ T cells, (C) CD8^+^ T cells, (D) CD4^+^ naive, effector, and active, and (E) CD8^+^ naive, effector, and active cells are shown. Data are pooled from two independent experiments (n = 9-10 mice/group). Data are expressed as mean ± SEM. Significant results were determined by two-way mixed ANOVA (* p ≤ 0.05, ** p ≤ 0.01, *** p ≤ 0.001, **** p ≤ 0.0001).

We similarly assessed CD8^+^ subpopulations including naïve, effector, and active cells. At day 4 p.i., RSV/PXQAD mice had a trend for reduced absolute cell number of effector cells and significant reductions in the absolute cell numbers for the naïve and active CD8^+^ subpopulations as compared to the RSV/PBS control mice (Fig. 4E and Table S1). In accordance, the percentage of effector and active CD8^+^ cells were decreased while the percent of naïve CD8^+^ cells increased (Table S1). At day 7 p.i., the absolute cell counts of effector and active CD8^+^ T-cells were significantly reduced (Fig. 4E). Similarly, the percent change of effector CD8^+^ cells was reduced, while the percent of naïve and active CD8^+^ T-cells remained comparable to the RSV/PBS control mice (Table S1). Collectively, these data indicate that HIF-1α plays an important role in regulating T-cell responses during RSV infection by modulating T-cell recruitment to the lung and their activation.

Last, we assessed the lung tissue of mice that received PXQAD by histopathology. Based on the scoring determined by a board-certified veterinarian with expertise in mouse lung morphology, uninfected mice that received PXQAD remained comparable to the PBS/PBS control mice in all categories and at all timepoints tested. RSV/PXQAD mice demonstrated significant improvements to the total lung score as compared to the RSV/PBS control mice (Fig. 5A). Specifically, perivasculitis (Fig. 5B) and peribronchiolitis (Fig. 5C) were significantly improved at days 4 and 7 p.i. as compared to the RSV/PBS control mice. This finding is consistent with the reduction of immune cells noted in the BAL and lung tissue of these mice. No significant differences in the percent of abnormal lung field (Fig. 5D) or the severity of interstitial pneumonia (Fig. 5E) were appreciated at any time point. Representative images of the lung tissue for RSV/PBS and RSV/PXQAD mice are shown in Fig. 6. Collectively, these data suggest HIF-1α to contribute to the worsening of airway function and tissue damage during the acute phase of RSV infection.

**Figure 5.**
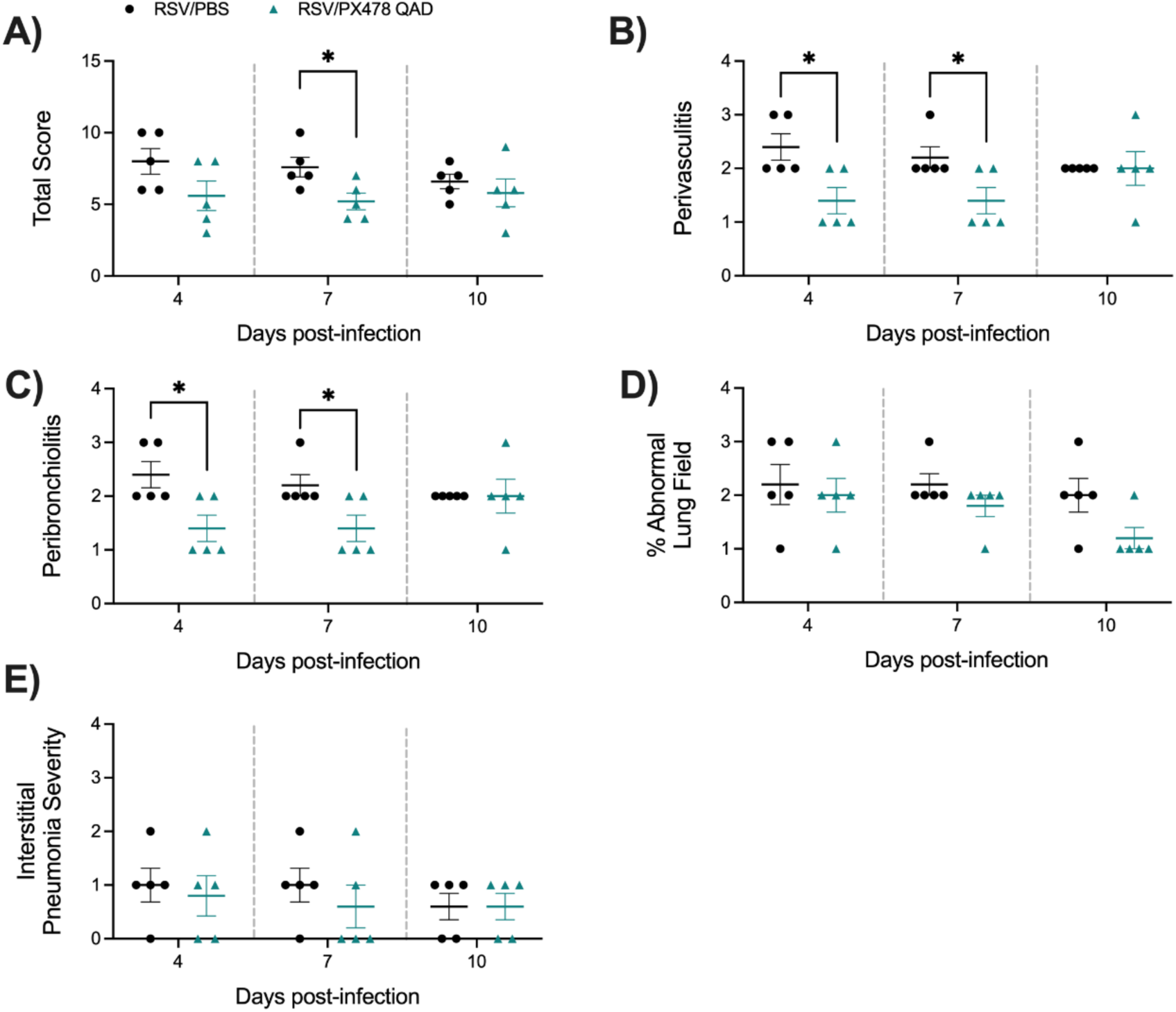
Histopathological scoring of lung tissue following HIF-1α inhibition during RSV infection. At days 4, 7, and 10 p.i., the left lung was collected from RSV infected mice that had received PBS or PX478 QAD and subjected to FFPE. Cuts of lung tissue were stained with H&E and observed under a microscope at 10X magnification. The (A) total score was calculated, consisting of scores for (B) perivasculitis, (C) peribronchiolitis, (D) percent abnormal lung field, and (E) interstitial pneumonia. Data are representative of one independent experiment (n = 5 mice/group). Data are expressed as mean ± SEM. Significant results were determined by an unpaired student’s t-test (* p ≤ 0.05).

**Figure 6.**
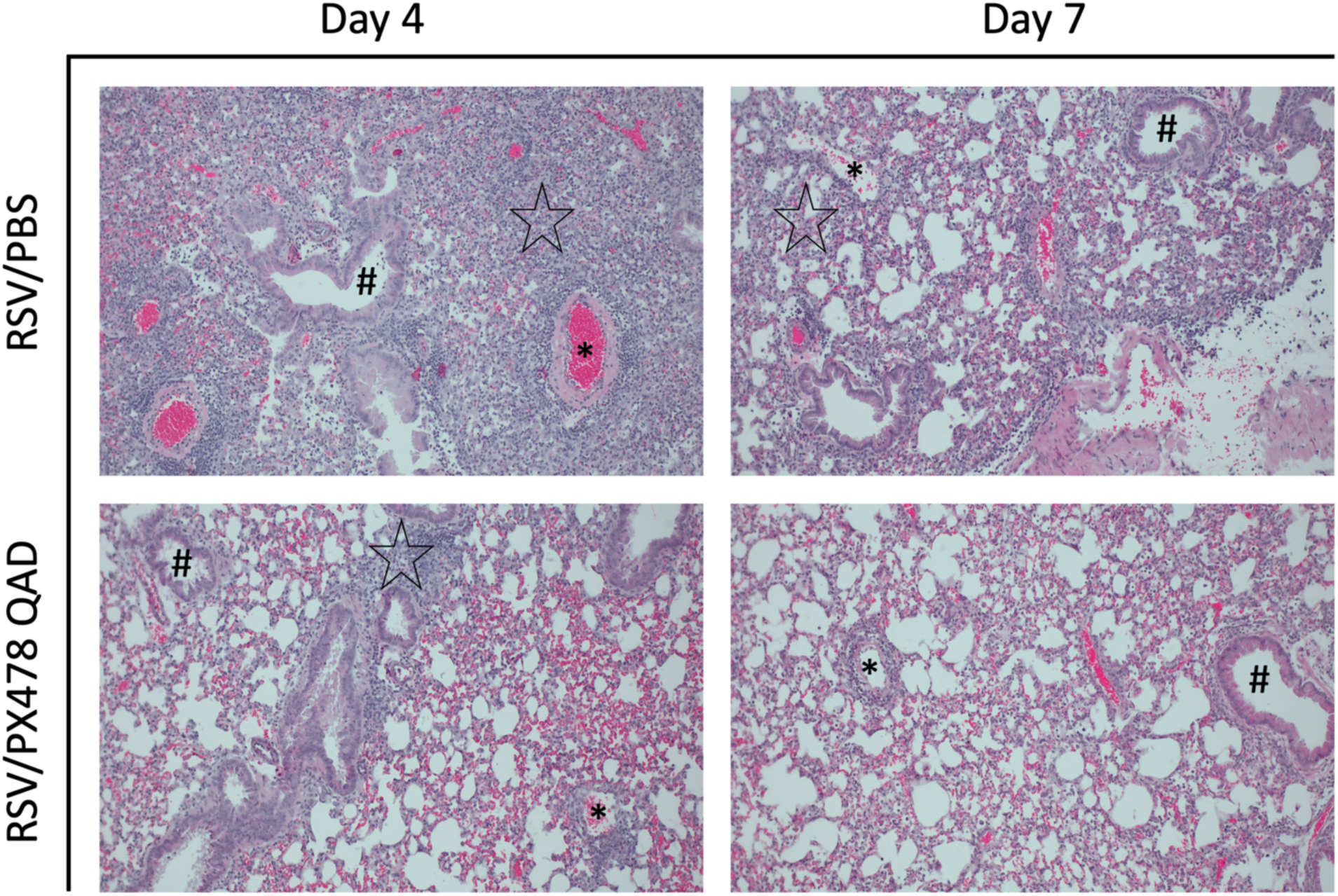
Histopathological imaging of lung tissue following HIF-1α inhibition during RSV infection. At days 4 and 7 p.i., the left lung was collected from mice that had received PBS or PX478 every 48 h and subjected to FFPE. Cuts of lung tissue were stained with H&E and observed under a microscope at 10X magnification. Perivasculitis is indicated by an asterisk. Peribronchiolitis is indicated by the hash sign. Interstitial pneumonia is indicated by the black star. No interstitial pneumonia present in RSV/PX478 QAD on day 7. Representative images are shown here (n = 5 mice/group).

### 4.4 Effect of PT2385 administration on RSV infection associated disease parameters and viral replication

While there is a degree of functional overlap between the HIF-α subunits, they have also been demonstrated to govern a distinct range of biological activities [14]. To better understand the specific role of HIF-2α during RSV infection, mice were administered PT2385 by oral gavage every 24 h at a dose of 50mg/kg (Fig. 7A). Shortly after the first dose, mice were anesthetized and infected with RSV at a dose of 5×10^6^ PFU. Uninfected mice that received PT2385 were found to have similar body weight loss and illness score as the PBS/CO control mice (Fig. 7B-7C). RSV infected mice that received PT2385 demonstrated significant improvements to bodyweight loss at several timepoints over the 10-day disease course as compared to the RSV/CO control mice (Fig. 7B). Interestingly, the illness score in RSV/PT2385 mice was worse at day 4 p.i., but treated animals showed a quicker recovery at later time points of infection when compared to the RSV/CO control mice (Fig. 7C).

**Figure 7.**
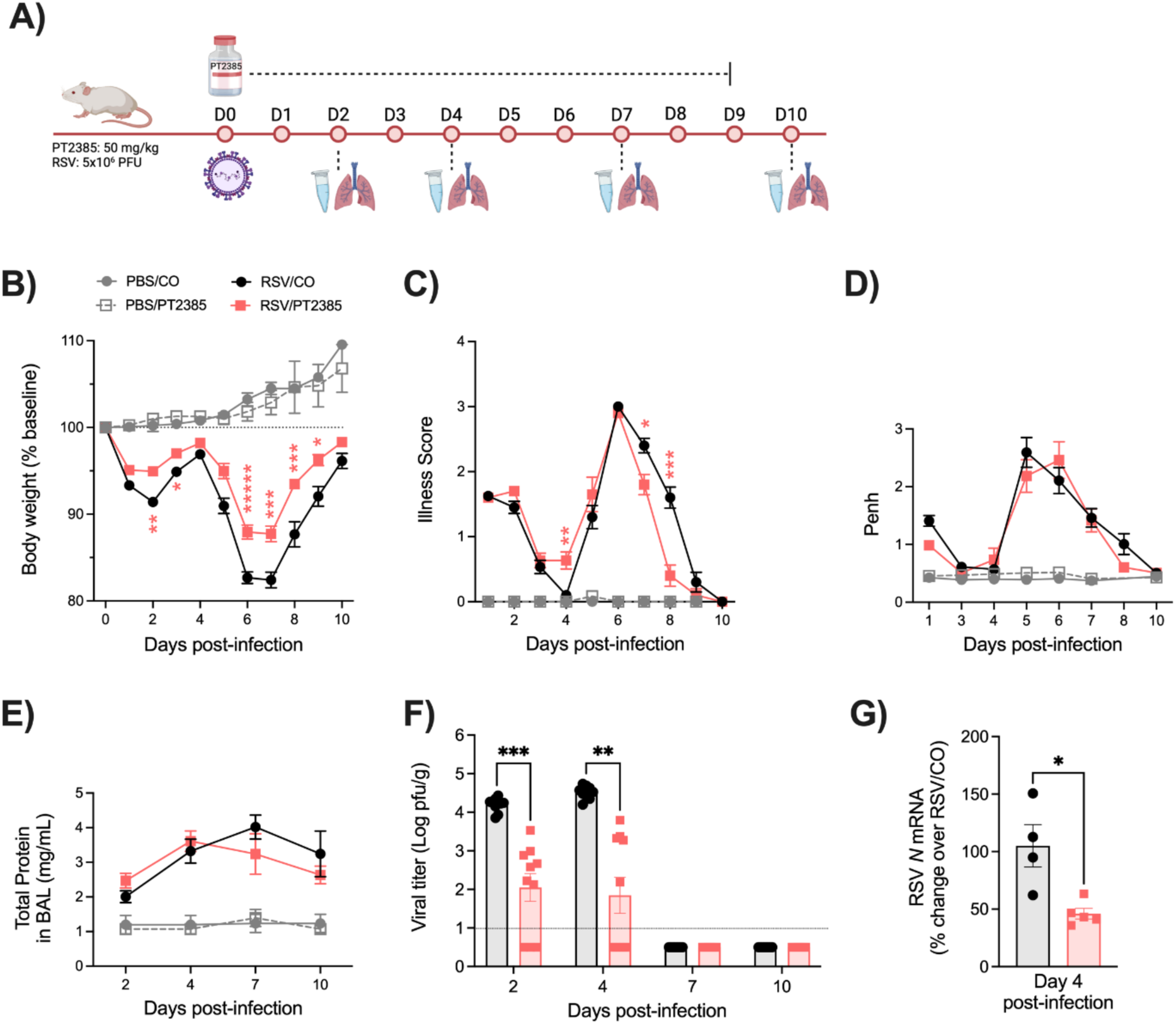
Assessment of clinical disease, airway function, and viral replication following HIF-2α inhibition during RSV infection. The experimental design for mice treated with PT2385 is described in (A). Following treatment, all mice were monitored daily for changes in (B) body weight and (C) illness score. (D) Bronchoconstriction, represented by baseline Penh, was measured by plethysmography. At days 2, 4, 7, and 10 p.i., (E) total protein was measured in the BAL fluid and the right lung was collected for assessment of (F) viral replication by plaque assay. At day 4 p.i, (G) RSV *N* mRNA levels were measured by RT-qPCR. For clinical disease, data are pooled from four independent experiments (PBS/CO n = 6-12, PBS/PT n = 6-18, RSV/CO n = 6-24, RSV/PT n = 6-24 mice/group). For bronchoconstriction, data are pooled from three independent experiments (PBS/CO and PBS/PT n = 6-12, RSV/CO and RSV/PT n = 6-24 mice/group). For total protein, data are pooled from two independent experiments (PBS/CO and PBS/PT n = 4-5, RSV/CO and RSV/PT n = 5-10 mice/group. For viral replication by plaque assay, data are pooled from two independent experiments (n = 9-10 mice/group). For RT-qPCR, data are from one independent experiment (n = 4-5 mice/group). Data are expressed as mean ± SEM. Significant results were determined by two-way mixed ANOVA (B-F) and unpaired t test (G) (* p ≤ 0.05, ** p ≤ 0.01, *** p ≤ 0.001, **** p ≤ 0.0001).

To assess airway function during RSV infection, we measured bronchoconstriction and BAL total protein levels. Uninfected mice that received PT2385 showed no significant alterations to Penh values at any timepoint tested as compared to the PBS/CO control mice. Similarly, RSV infected mice treated with PT2385 had similar Penh values as the RSV/CO control mice throughout the 10-day disease course (Fig. D). We observed no significant differences in total protein for all mice tested as compared to their respective control groups (Fig. 7E).

Assessment of viral replication in the lung tissue, assessed by plaque assay and viral gene expression, showed RSV/PT2385 mice to have reduced RSV replication at early and peak timepoints of days 2 and 4 p.i. (Fig. 7F and 7G). By days 7 and 10 p.i., no replicating virus was detected in RSV/PT2385 mice, indicating successful clearance of RSV from the lung tissue. Collectively, these data suggest HIF-2α contributes to RSV-induced disease and control of viral replication.

### 4.5 Effect of PT2385 administration on RSV-induced HIF-2α target genes

To confirm inhibition of RSV-induced HIF-2α activation by PT2385 treatment, we analyze the expression of *Epas1* (HIF-2α) and the HIF-2α-target genes *Serpine1* and *Epo* at day 2 p.i. in lungs of mice untreated or treated with the compound (Fig. 8). There was no change in baseline HIF-2α gene expression (*Epas1*) following RSV infection, different from what we observed for HIF-1α. We confirmed induction of the HIF-2α-target gene *Serpine1*, which was significantly reduced by PT2385 treatment. *Epo* was not detectible in the lung tissue of any mice. There was no change in RSV-induced expression of HIF-1α-dependent genes *Hk2, Slc2a1*, and *Pgk1* following treatment, indicating specificity of the compound (Fig. 8).

**Figure 8.**
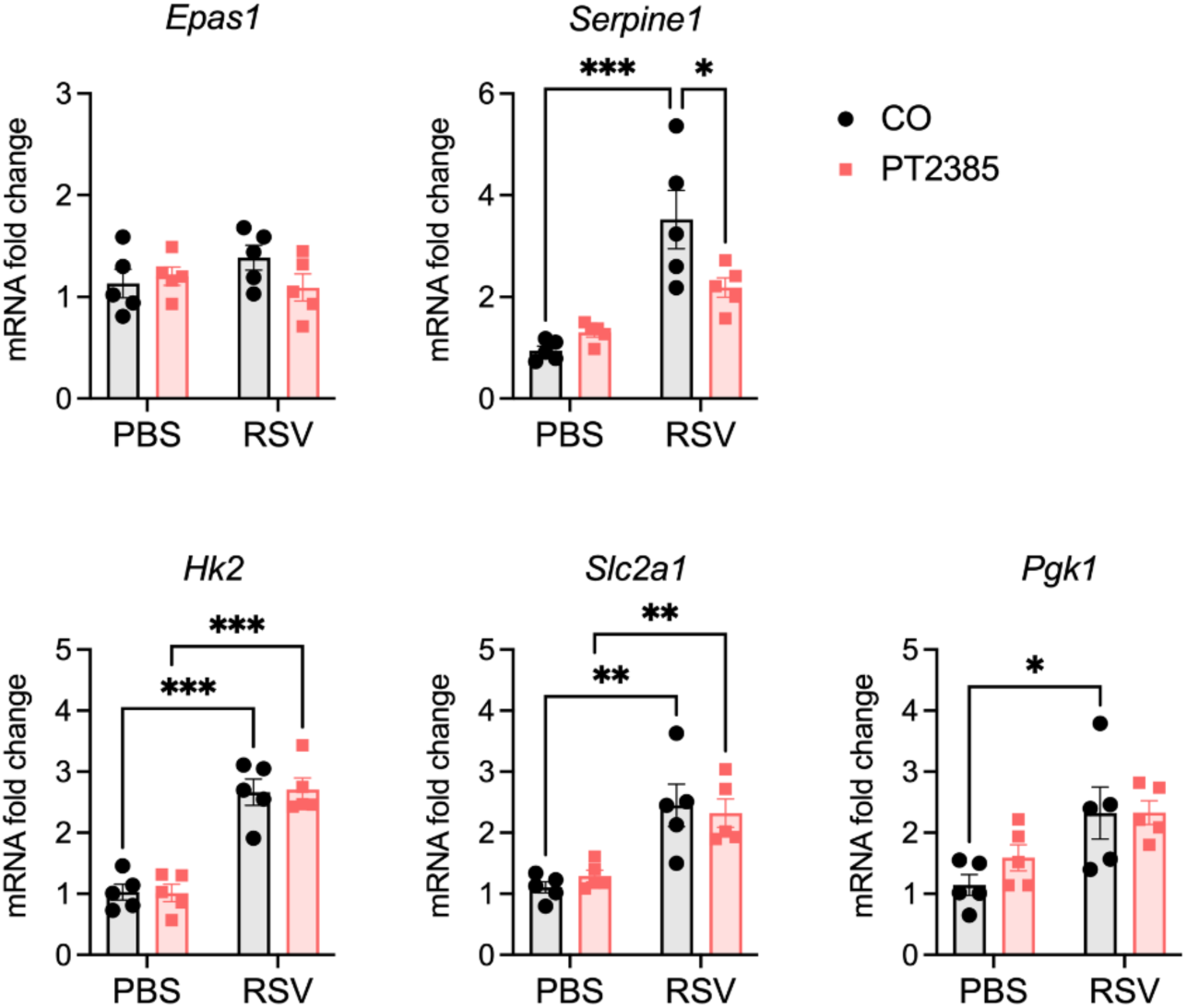
Effect of PT2385 on Epas1 (HIF-2α) and HIF-1α and -2α target genes in lungs of mice during RSV infection. A group of mice were euthanized at day 2 p.i., lungs were harvested, and total lung RNA was extracted. *Epas1* (HIF-2α), the HIF-2α target *Serpine1* (PAI-1), and the HIF-1α targets *Pgk1, Slc2a1* (GLUT1) and *Hk2* gene expression was measured by RT-qPCR (n = 5 mice/group). Data are expressed as mean ± SEM. Significant results were determined by two-way ANOVA followed by Bonferroni’s multiple comparisons test (* p ≤ 0.05, ** p ≤ 0.01, *** p ≤ 0.001).

### 4.6 Inhibition of HIF-2α leads to increased immune activity in the BAL and lung tissue of RSV infected mice

We had previously demonstrated HIF-2α to be dispensable for the control of RSV replication in human small airway epithelial cell [23]. Given the antiviral outcome in RSV/PT2385 mice, we next wanted to understand the contribution of HIF-2α to the immune response during an RSV infection. We first evaluated the cellular composition of the BAL fluid. Uninfected mice that received PT2385 showed no significant difference in BAL cellular composition, compared to the PBS/CO control group (Fig. 9A-D). RSV infected mice that received PT2385 demonstrated a significant increase in the number of total of cells present in the BAL throughout the course of the infection (Fig. 9A), particularly for the number of lymphocytes (Fig. 9B), compared to the PBS/CO control group. Macrophage cell counts were also significantly increased at days 2 and 10 p.i. (Fig. 9C), while neutrophil cell counts remained comparable to the RSV/CO control mice (Fig. 9D).

**Figure 9.**
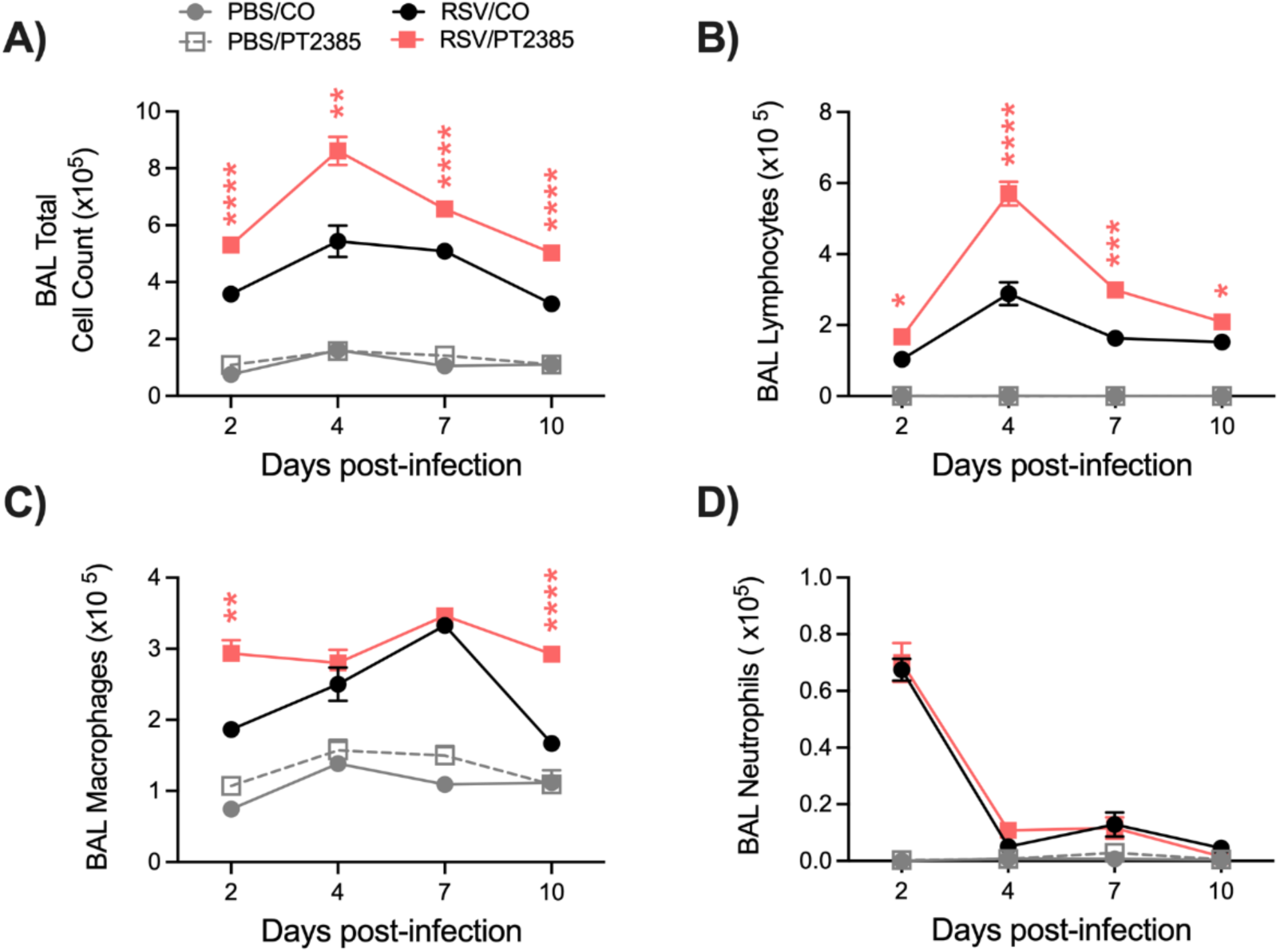
Differential cell count of the BAL fluid following HIF-2α inhibition during RSV infection. At days 2, 4, 7, and 10 p.i., BAL fluid was collected and used to obtain (A) total cell counts, as well as differential cell counts consisting of (B) lymphocytes, (C) macrophages, and (D) neutrophils. Data are pooled from two independent experiments (PBS/CO and PBS/PT n = 4-6, RSV/CO and RSV/PT n = 10-15 mice/group). Data are expressed as mean ± SEM. Significant results were determined by mixed-effects analysis (* p ≤ 0.05, ** p ≤ 0.01, *** p ≤ 0.001, **** p ≤ 0.0001).

Given the increase in some innate cells in the BAL, we next assessed a few innate immune cells speculated to contribute to RSV antiviral activity by flow cytometry. Whole lung tissue was collected from RSV infected mice treated or not with PT2385 at days 1 and 2 p.i., the peak of innate immune cell activity in RSV infected mice. We assessed the tissue for alveolar macrophages, neutrophils, natural killer (NK), and natural killer T (NKT) cells. We found a significant increase in absolute cell count of neutrophils on day 2 p.i. in the lungs of RSV/PT2385 mice as compared to the RSV/CO control mice (Fig. 10A). Absolute cell counts of alveolar macrophages, NK, and NKT cells were comparable between the two groups (Fig. 10B-D).

**Figure 10.**
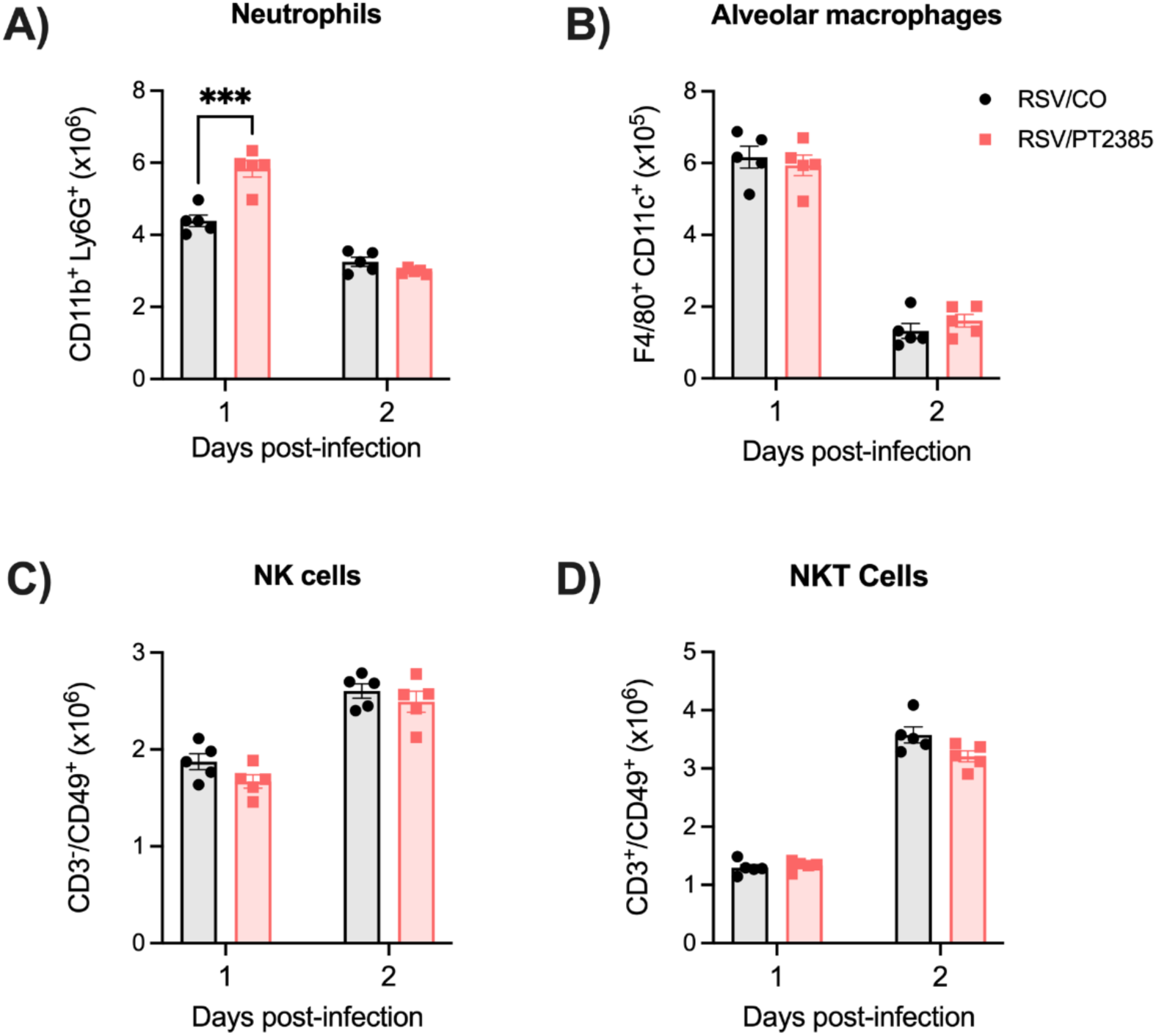
Assessment of neutrophils, alveolar macrophages, natural killer, and NKT cells following HIF-2α inhibition during RSV infection. Mice were euthanized at days 1 and 2 p.i., and whole lung tissue was collected. Single cell suspension was prepared, stained with live/dead cell dye and fluorochrome-conjugated antibodies, and acquired by flow cytometry. Data were analyzed following the gating strategy in Figure S3. Shown here are absolute cell counts for (A) neutrophils, (B) alveolar macrophages, (C) NK cells, and (D) NKT cells. Data are expressed as mean ± SEM (n = 5 mice/group). Significant results were determined by two-way ANOVA (*** p ≤ 0.001).

As with HIF-1α, we next assessed CD4^+^ and CD8^+^ T-cell responses by flow cytometry analysis. RSV infected mice treated with PT2385 demonstrated a significant increase of total leukocytes (Fig. 11A), CD4^+^ (Fig. 11B) and CD8^+^ (Fig. 11C) T-cells in the lung at day 4 p.i.. We further analyzed the CD4^+^ T-cell subpopulations of naïve, effector, and active T-cells. In RSV/PT2385 mice at day 4 p.i., we observed an increasing trend in total number of CD4^+^ naïve cells as well as a significant increase in the absolute number of CD4^+^ effector and active T-cells as compared to the RSV/CO control mice (Fig. 11D). The percent change at day 4 p.i. was similar amongst all CD4^+^ T-cell subpopulations when compared to the RSV/CO control mice (Table S2). By day 7 p.i., the absolute cell counts and percent change of all CD4^+^ T-cells in the RSV/PT2385 were of comparable number to the RSV/CO control mice. For CD8^+^ T-cell subpopulations, we observed a significant increase in absolute number of CD8^+^ naïve cells at day 4 p.i., while the absolute cell counts for effector and active CD8^+^ T-cells remained similar to the RSV/CO control mice (Fig. 11E). At day 7 p.i., the absolute number of active CD8^+^ T-cells was significantly increased, while the naïve and effector CD8^+^ cells remained comparable to the RSV/CO control mice (Fig. 11E). The percent change was not modified for any CD8^+^ cells at either timepoint (Table S2). Collectively, these data show that inhibition of HIF-2α results in increased T-cell numbers, which could contribute to better control of RSV replication, resulting in decreased viral titers.

**Figure 11.**
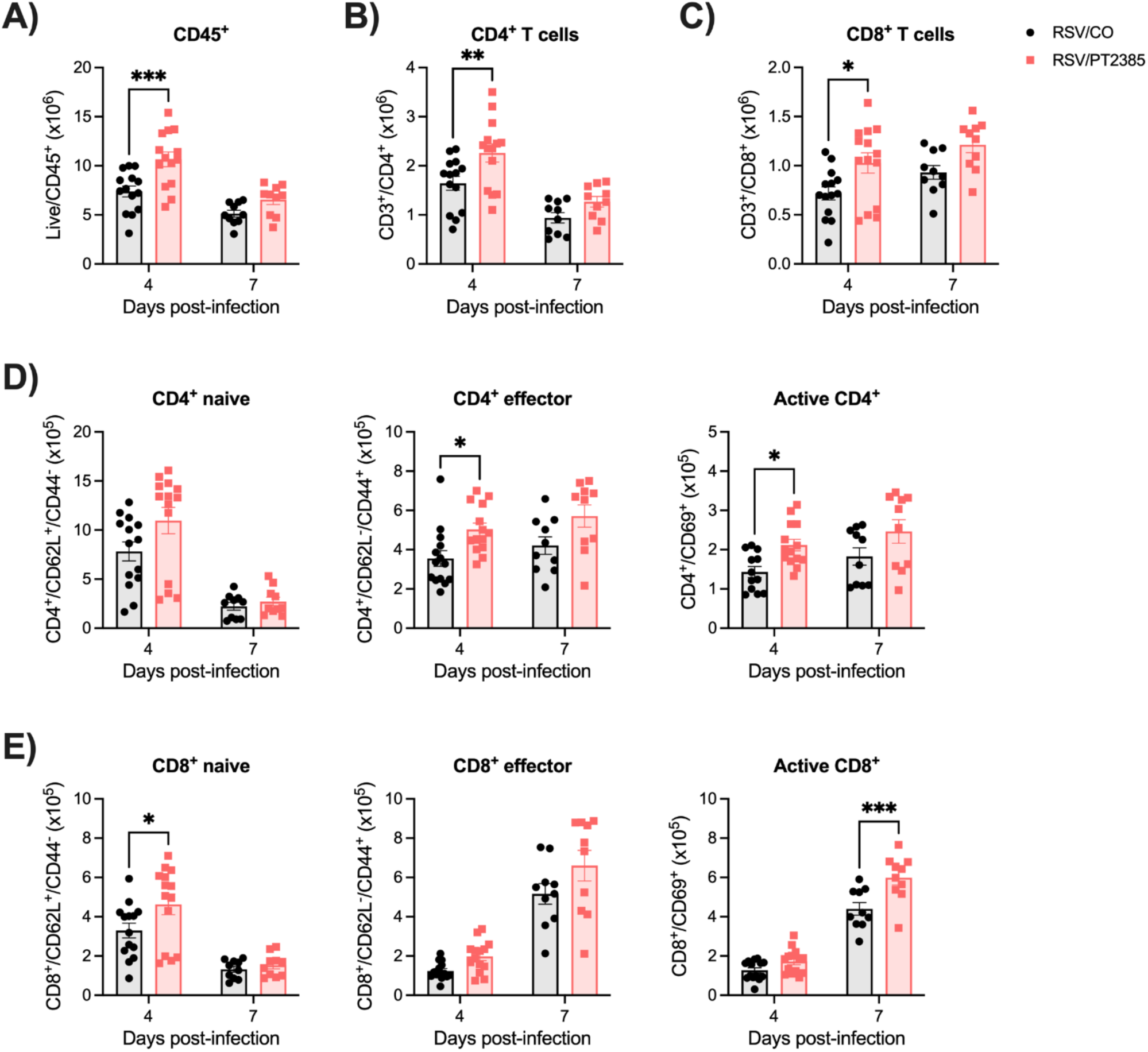
Assessment of CD4^+^ and CD8^+^ T-cells following HIF-2α inhibition during RSV infection. Whole lung tissue was collected at peak RSV replication and a timepoint of viral clearance, days 4 and 7 p.i., respectively. Single cell suspension was prepared, stained with live/dead cell dye and fluorochrome-conjugated antibodies, and acquired by flow cytometry. Data were analyzed following the gating strategy in Figure S2. Quantification of absolute cell counts of (A) Live CD45^+^, (B) CD4^+^ T cells, (C) CD8^+^ T cells, (D) CD4^+^ naive, effector, and active, and (E) CD8^+^ naive, effector, and active cells are shown. Data are pooled from two to three independent experiments (n = 10-14 mice/group). Data are expressed as mean ± SEM. Significant results were determined by a two-way mixed ANOVA (* p ≤ 0.05, ** p ≤ 0.01, *** p ≤ 0.001).

Lastly, we assessed the lung tissue of mice that received PT2385 by histopathology. Uninfected mice that received PT2385 remained comparable to the PBS/CO control mice in all categories and at all time points tested. RSV infected mice treated with PT2385 demonstrated significant improvements at day 10 p.i. in total lung score (Fig. 12A), peribronchiolitis (Fig. 12C), and the severity of interstitial pneumonia (Fig. 12E) as compared to the RSV/CO control mice. The percent abnormal lung field was found to have a trend towards improvement in RSV/PT2385 mice (Fig. 12D). Additionally, there was mild worsening of perivasculitis compared to the RSV/CO control mice (Fig. 12B). Representative images of the lung tissue for RSV/CO and RSV/PT2385 mice are shown in Fig.13. Collectively, these data suggest HIF-2α does not modify airway function but does contribute to worsening lung pathology during the acute phase of RSV infection.

**Figure 12.**
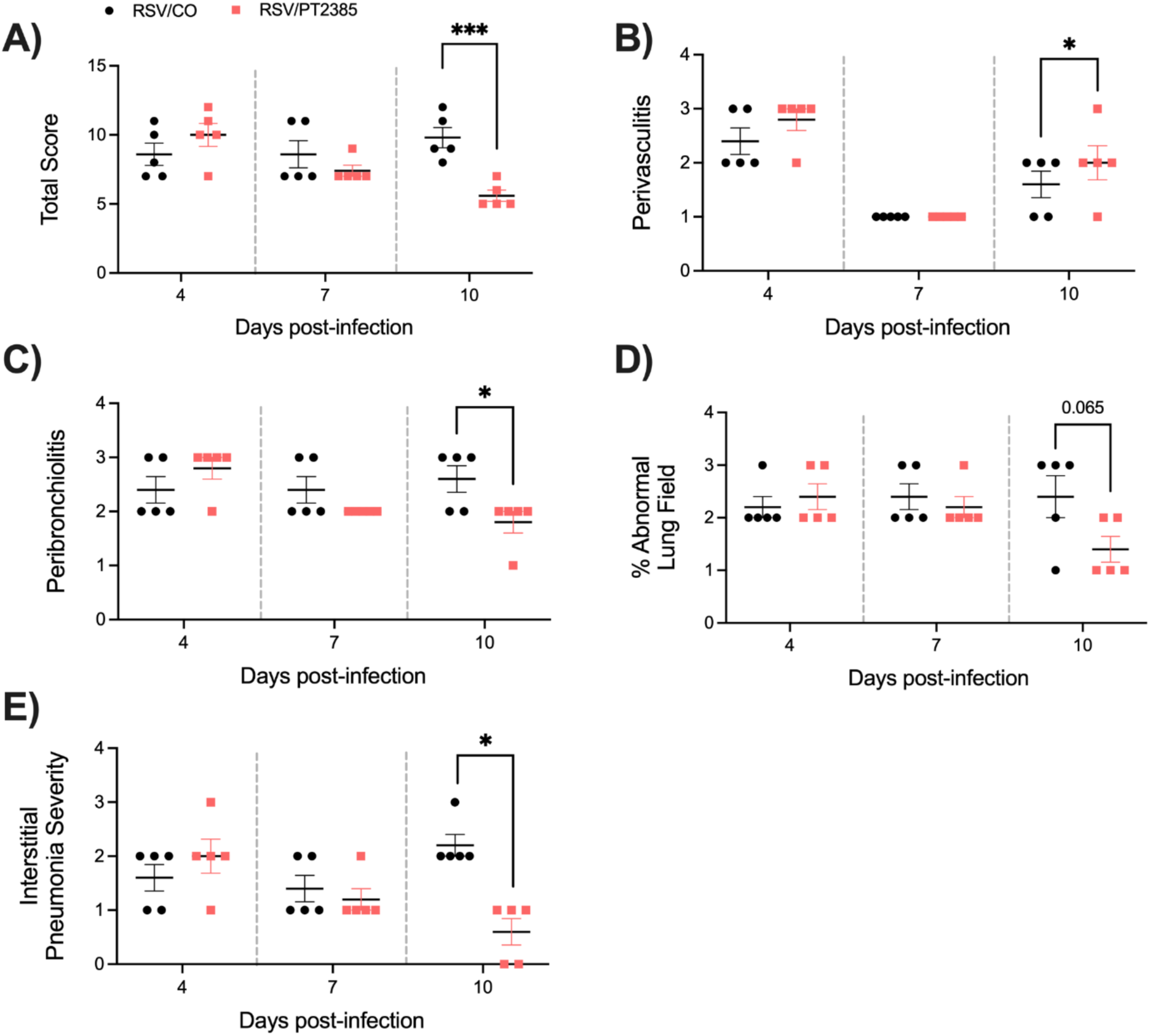
Histopathological scoring of lung tissue following HIF-2α inhibition during RSV infection. At days 4, 7, and 10 p.i., the left lung was collected from RSV infected mice that had received CO or PT2385 and subjected to FFPE. Cuts of lung tissue were stained with H&E and observed under a microscope at 10X magnification. The (A) total score was calculated, consisting of scores for (B) perivasculitis, (C) peribronchiolitis, (D) percent abnormal lung field, and (E) interstitial pneumonia. Data are representative of one independent experiment (n = 5 mice/group). Data are expressed as mean ± SEM. Significant results were determined by an unpaired student’s t-test (* p ≤ 0.05, *** p ≤ 0.001).

**Figure 13.**
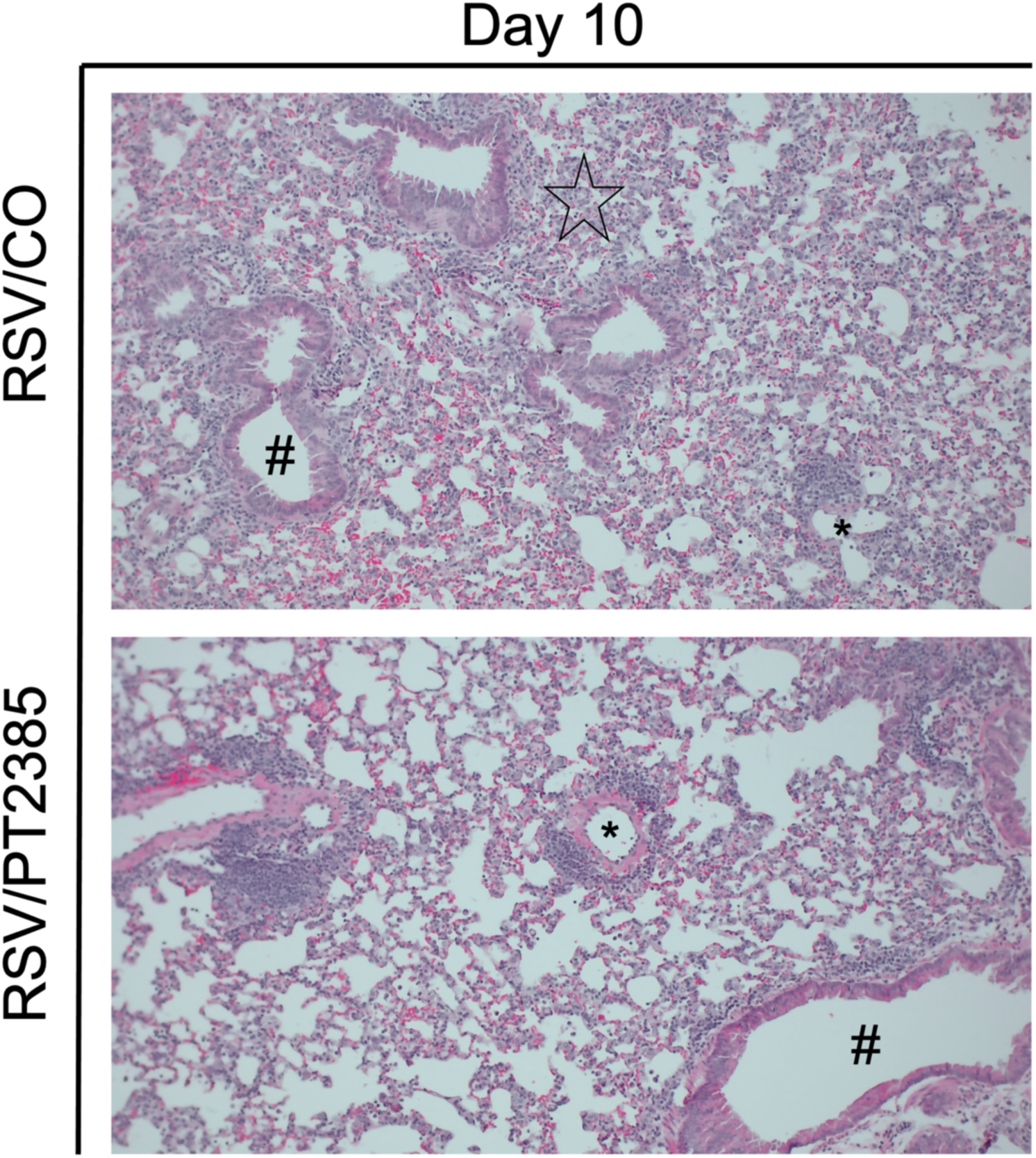
Histopathological images of lung tissue following HIF-2a inhibition during RSV infection. At day 10 p.i., the left lung was collected from mice that had received PBS or PT2385 and subjected to FFPE. Cuts of lung tissue were stained with H&E and observed under a microscope at 10X magnification. Perivasculitis is indicated by an asterisk. Peribronchiolitis is indicated by the hash symbol. Interstitial pneumonia is indicated by the black star. No interstitial pneumonia present in RSV/PT2385. Representative images are shown here (n = 5 mice/group).

## 5 Discussion

The pharmacological use of HIF therapeutics has been extensively explored in various models of cancer, autoimmunity, and non-infectious lung damage [7, 22, 23]. While some studies have demonstrated the antiviral potential of HIF inhibitors against respiratory viruses, these findings have been primarily limited to *in vitro* characterizations [8]. Our laboratory has previously shown that RSV infection results in the stabilization of both HIF-1α and HIF-2α. Through pharmacological inhibition of HIF, we also demonstrate the stabilization of HIF-1α, but not HIF-2α, to orchestrate metabolic reprogramming towards glycolysis and for this activity to be important for RSV replication in lung epithelial cells [14]. Given the diverse biological role of HIFs across several cellular compartments, we next wanted to understand the function of these inhibitors in a mouse model of RSV infection. In this study, we aimed to characterize the disease modulating properties and antiviral potential of HIF-1α and -2α inhibitors in a BALB/c mouse model of RSV infection.

Our results show that inhibition of HIF-1α worsens clinical disease while concurrently improving aspects of airway function and lung inflammation during RSV infection. In our initial experiments with PX478, when administering the inhibitor every-day (QD), we observed notable weight loss and increased illness in the PBS/PXQD mice compared to the PBS control mice. As there was no difference in any other parameter analyzed, including BAL cells (data not shown), this result suggests that the deteriorating clinical condition in PBS/PXQD mice is likely influenced by a factor other than inflammation or tissue damage. Previous studies investigating the anti-tumor efficacy of PX478 have administered this compound for up to 5 consecutive days, reporting minor weight-loss in the mice during that time [24, 25]. Studies using obese mouse models have noted PX478 to prevent weight gain [26, 27] and a recent investigation has shown the efficacy of PX478 as a potential therapeutic for type-II diabetes [28]. Considering the significant role of HIF-1α in metabolic function and the recent repositioning of PX478 for treatment of type-II diabetes, it is possible that consecutive dosing of PX478 for extended periods of time may lead to appetite suppression. In our model, spacing treatments over 48 h (QAD) ameliorated this effect, resulting in patterns of clinical disease in the PBS/PXQAD mice that resemble previously reported observations. Despite this, RSV infected mice treated with PX478 every 48h continued to display delayed recovery as compared to the RSV control mice (Fig. 1F). This was surprising as these RSV infected mice effectively cleared the virus from the lung tissue and show no signs of increased inflammatory activity. One possible explanation for the phenomenon could be a moderate appetite suppression induced by PX478 that is amplified due to the stress of infection.

Treatment with PX478 in RSV infected mice resulted in initial antiviral activity regardless of the PX478 dosing schedule. This was followed by prolonged viral replication at later time points of infection when PX478 was given QD, but not QAD. Concurrently, these mice experienced a significant reduction in the number of total cells present in the BAL, more strikingly in the number of lymphocytes. HIF-1α is critical for the activation, migration, and cytotoxic functions of T-cells [29]. T-cells play an important role in controlling RSV replication and clearance within the host [20–22]. During the early stages of RSV disease, CD4^+^ effector cells are known to aid in the recruitment and activation of other immune cells that control RSV replication such as natural killer (NK) cells, CD8^+^ cytotoxic T-cells, and B-cells [30–32]. At later stages of disease, CD8^+^ effector T-cells are responsible for directly clearing RSV from the lung epithelium [33]. RSV/PXQAD mice had a significant reduction of total leukocytes, total CD4^+^ T-cells, and total CD8^+^ T-cells at both timepoints assessed. Furthermore, during peak of viral replication, the absolute and percentage of active and effector CD4^+^ T-cells were significantly reduced. Similarly, PXQAD treatment resulted in significantly reduced number of active and effector CD8^+^ T-cells present during peak viral clearance in the lungs of infected mice. Despite the overall impact on the T-cell population, the RSV/PXQD and RSV/PXQAD treated mice displayed antiviral activity at early and/or peak timepoints of RSV replication. Given the suppressed T-cell activity, this antiviral effect is more likely attributed to impaired RSV replication rather than immune-mediated activity. Aligning with our previous *in vitro* findings, the current study shows that HIF-1α inhibition in RSV-infected mice influences the expression of genes that regulate critical metabolic pathways, including glycolysis, an essential pathway for RSV replication in airway epithelial cells [15]. This likely contributes to the reduced RSV replication at early and peak timepoints in mice, rather than immune-mediated activity.

Several studies have shown CD8^+^ T-cells to be responsible for the clearance of RSV from the lung tissue [33]. In our PXQD model, RSV replication is prolonged, indicating that the amount of virus that can replicate is not being cleared from the lung tissue (Fig S1). CD8^+^ T-cells were found to be further suppressed in the RSV/PXQD mice when compared to the RSV/PXQAD mice (data not shown). These results would suggest continuous treatment with PX478 to result in a threshold effect of CD8^+^ T-cell suppression, preventing efficient viral clearance. This is an important finding, considering the use of HIF modulating therapeutics in the context of other diseases, such as cancer.

In contrast to HIF-1α, inhibition of HIF-2α by PT2385 was found to improve clinical disease, with no difference in airway function. Like PX478, treatment with PT2385 demonstrated significant antiviral activity at early and peak timepoints of RSV replication, with no impact on viral clearance. Based on our previous characterizations in epithelial cells, where inhibition of HIF-2α did not affect metabolic reprogramming and viral replication [15], the mechanisms underlying this activity could be distinct from the ones responsible for impaired replication following HIF-1α inhibition.

RSV/PT2385 mice displayed a moderate increase in immune cell presence in the airway space. This was characterized by an increased number of BAL macrophages and lymphocytes as well as lung neutrophils and active CD4^+^ and CD8^+^ T-cells. Neutrophilic responses have been shown to contribute to the control of viral replication in several models of respiratory virus infection, including RSV, influenza, and SARS-CoV-2 [34–37], although other unidentified mechanism(s) are likely to play an important role in the early antiviral activity observed with PT2385 administration. Recent studies found that a loss of HIF-2α in regulatory T-cells (Treg) was associated with reduced suppressor functions, leading to an increase in CD4^+^ T-cell activity, without changes in Treg development [38, 39]. In agreement with this finding, our results indicate an increase in CD4^+^ and CD8^+^ T-cell number, including active/effector cells, with no significant alteration to Treg cell numbers (data not shown), compared to the RSV/PBS control mice. As for other models of infection/inflammation, loss of HIF protein activation in myeloid cells in a model of Leishmania infection also resulted in heightened CD4^+^ Th1 immune response [40], and mice with T-cell intrinsic HIF-2α deletion showed exacerbated intestinal inflammation, in a model of experimental colitis [41]. Since HIF-2α was shown to be dispensable for RSV replication in epithelial cells, it is possible that this increase in immune cell responses, both innate and adaptive, could play a major role in the observed antiviral effect seen in RSV/PT2385 mice.

HIF-1α activation has been demonstrated to impact airway function in a variety of ways, including the induction of bronchoconstriction [42]. HIF-1α achieves this by influencing the contractility of airway smooth muscles through several mechanisms such as stimulating calcium production, modulating ion channels, and triggering the release of mediators like leukotrienes [43–45]. In contrast, HIF-2α has been shown to be dispensable for the induction of changes to airway hypersensitivity and bronchoconstriction [38]. Consistent with these findings, we observed notable improvements in bronchoconstriction in RSV-infected mice treated with PX478 but not with PT2385. In the PXQAD experiments, pathology scores for perivasculitis and peribronchiolitis were significantly improved on days 4 and 7 p.i., as compared to the RSV control mice. This finding is consistent with the reduced immune cell infiltration and antiviral activity at these timepoints. As HIF-1α has important roles at the later stages of lung injury in promoting angiogenesis and tissue regeneration [46], it is possible that inhibition of HIF-1α at the later stages of disease may hinder proper tissue repair, negating the benefit observed in the initial phase of RSV lung disease. Mice treated with PT2385 during RSV infection had no major alterations to any scored histopathology category until day 10 p.i., despite the observed increase in BAL cellularity and increased T-cell recruitment to the lung, which could possibly explain the lack of improvement in airway function of these mice. By the end of the acute phase, RSV/PT2385 mice had a mild increase in perivasculitis, but significant improvements in all other categories. This increase in perivasculitis could be due to the increase in the number of immune cells observed within the BAL and lung tissue, but could also be related to changes in vascular function, as HIF-2α plays an important role in maintaining the vascular integrity [47].

With the growing appreciation for the viral manipulation of the HIF pathway and the targeted therapeutic potential, several studies have begun to investigate the contribution of HIF proteins to diseases associated with other respiratory viruses. SARS-CoV-2 infection was found to induce stabilization of HIF-1α in immune cells, via mitochondrial damage, facilitating viral replication and cytokine production [48]. On the other hand, HIF activation via the prolyl hydroxylase inhibitor Roxadustat reduced angiotensin-converting enzyme 2 (ACE2) expression and inhibited SARS-CoV-2 entry and replication both in lung epithelial cells [49], as well as in a hamster model of infection [50]. *In vitro*, IAV infection induces HIF stabilization via inhibition of the proteosome and decreased expression of factor-inhibiting-HIF (FIH), with no changes in HIF-1α mRNA transcription or hydroxylation [51], with pharmacological inhibition of HIF significantly reducing viral replication [52]. In contrast, HIF-1α deficiency in lung epithelium was found to facilitate viral replication in a mouse model of influenza A virus infection by promoting autophagy [53].

Collectively, our data demonstrate HIF-1α and HIF-2α to be important regulators of the immune response to RSV infection. We also highlight HIF-1α as a critical component to the onset of RSV-induced bronchoconstriction. Additionally, these data describe the novel and complex contribution of the HIF-α subunits to RSV replication and clearance. Though we show antiviral potential for both HIF-α inhibitors, the resulting impact to clinical disease and modulation of the immune responses warrant great care be taken when considering HIF-α inhibitors for therapeutic use against a viral respiratory infection. Furthermore, our data demonstrates that the continued inhibition of HIF-1α can result in prolonged viral replication, highlighting the possible unwanted side effects associated with the use of HIF modulating therapeutics in the context of other diseases, such as cancer.

Future investigations will focus on further understanding the mechanistic contribution of the different HIF alpha subunits to the immune response and tissue repair processes during RSV infection.

## 7 Author Contributions

Performed experiments, D.R.M., Y.Q., T.I., M.W., A.H.M., and T.L; performed histopathology analysis, Y.L.J.H; contributed to data analysis and drafting of the manuscript, D.R.M. and A.H.M.; contributed to the conception of the manuscript, literature review, and final manuscript editing, D.R.M., A.C. and R.P.G. All authors have read and agreed to the published version of the manuscript.

## 9 Acknowledgments

We would like to thank Vineet Menachery, Yuejin Liang, Casey Wright, and Xiaoyi Yuan for their thoughtful discussion. We would also like to thank the Texas A&M University Veterinary Medicine and Biomedical Sciences Research Histology Core Facility for their assistance with the tissue processing. Biorender.com was used to produce the experimental design summaries.

## 10 Supplementary Material

**Figure S1.**
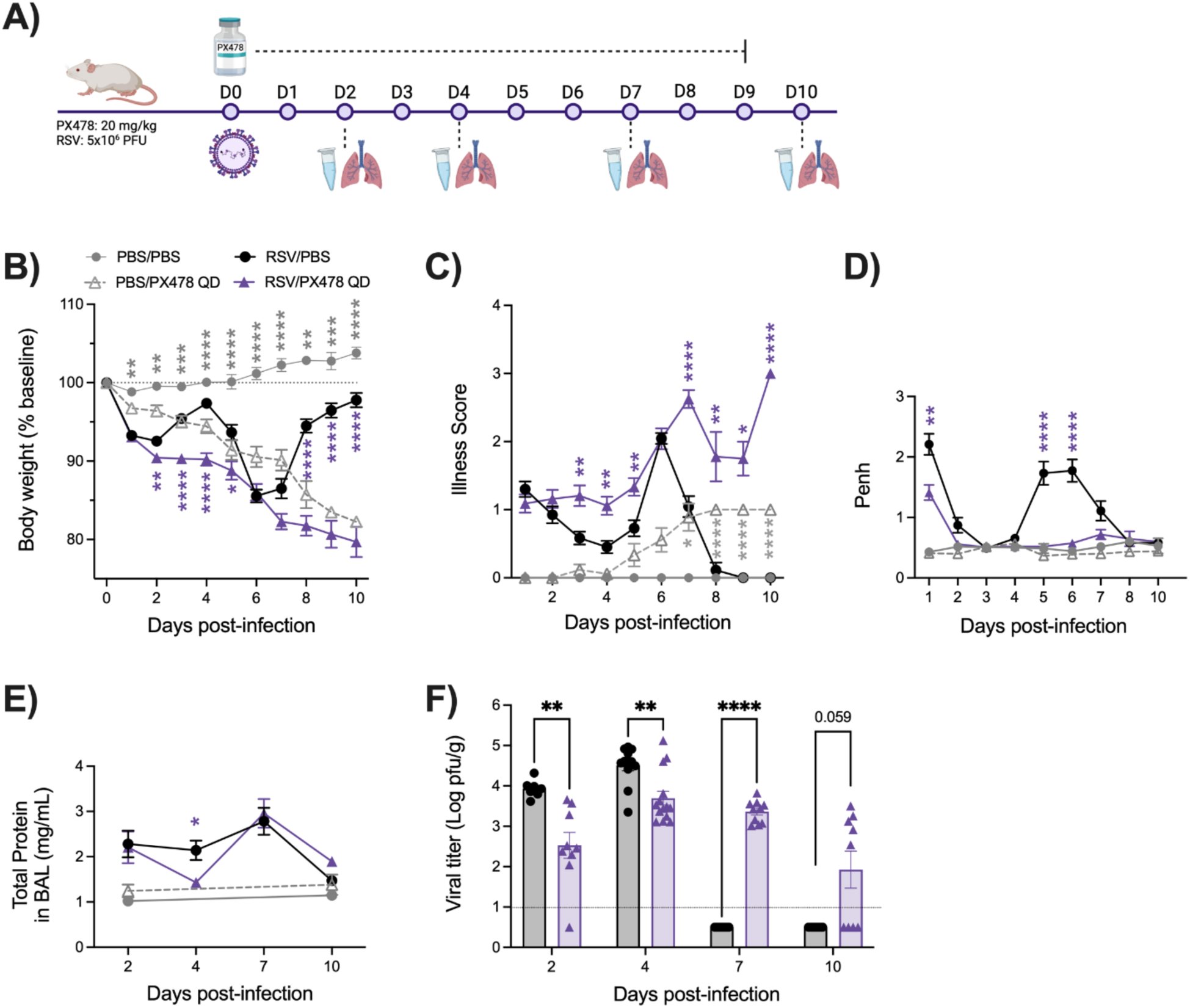
Assessment of clinical disease, airway function, and viral replication following HIF-1a inhibition during RSV infection. The experimental design for mice treated with PX478 QD is described in (A). Following treatment, all mice were monitored daily for changes in (B) body weight and (C) illness score. (D) Bronchoconstriction, represented by baseline Penh, was measured by plethysmography. At days 2, 4, 7, and 10 p.i., (E) total protein was measured in the bronchoalveolar lavage (BAL) fluid and the right lung was collected for assessment of (F) viral replication by plaque assay. For clinical disease, data are pooled from four independent experiments (PBS/PBS n = 6, PBS/PX n = 6-18, RSV/PBS and RSV/PX n = 6-24 mice/group). For bronchoconstriction, data are pooled from three independent experiments (PBS/PBS and PBS/PX n = 6, RSV/PBS and RSV/PX n = 6-24 mice/group). For total protein, data are pooled from two independent experiments (PBS/PBS and PBS/PX n = 4, RSV/PBS and RSV/PX n = 8-10 mice/group). For viral replication, data are pooled from two independent experiments (RSV/PBS n = 7-12, RSV/PX n = 9-14 mice/group). Data are expressed as mean ± SEM. Significant results were determined by two-way mixed ANOVA (* p ≤ 0.05, ** p ≤ 0.01, *** p ≤ 0.001, **** p ≤ 0.0001).

**Figure S2.**
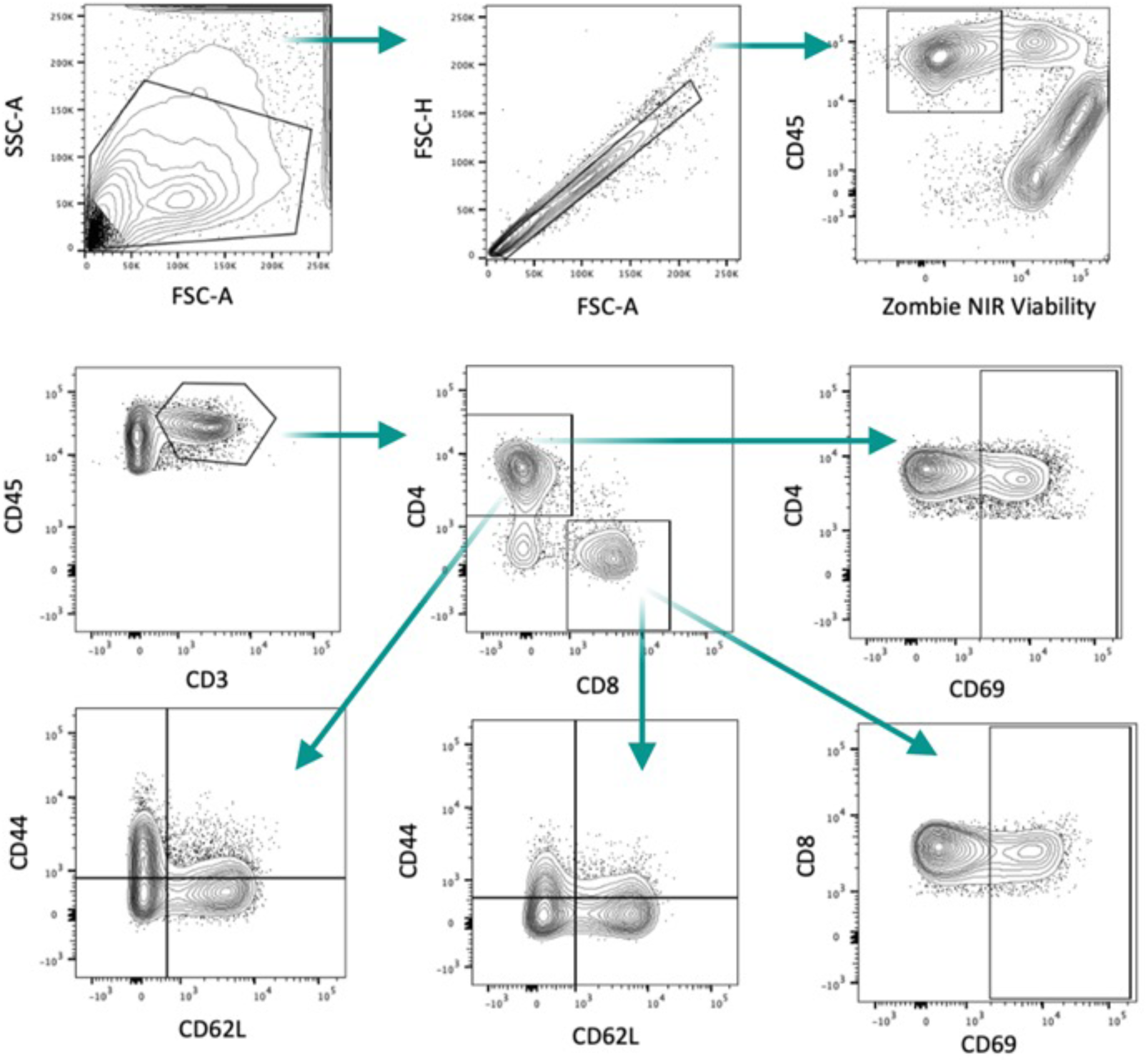
Schematic for analysis of CD4^+^ and CD8^+^ T-cells by flow cytometry. At days 4 and 7 p.i., whole lung tissue was collected from the respective groups. Single cell suspension was prepared, stained with live/dead cell dye and fluorochrome-conjugated antibodies, followed by flow cytometric analysis. T cells were gated on CD45^+^CD3^+^ first, then further gated on CD4^+^ and CD8^+^ subpopulations. Naïve and effector T cells were gated on CD44^−^CD62L^+^ and CD44^+^CD62L^−^, respectively. CD69 was used as an early activation marker of T cells.

**Figure S3.**
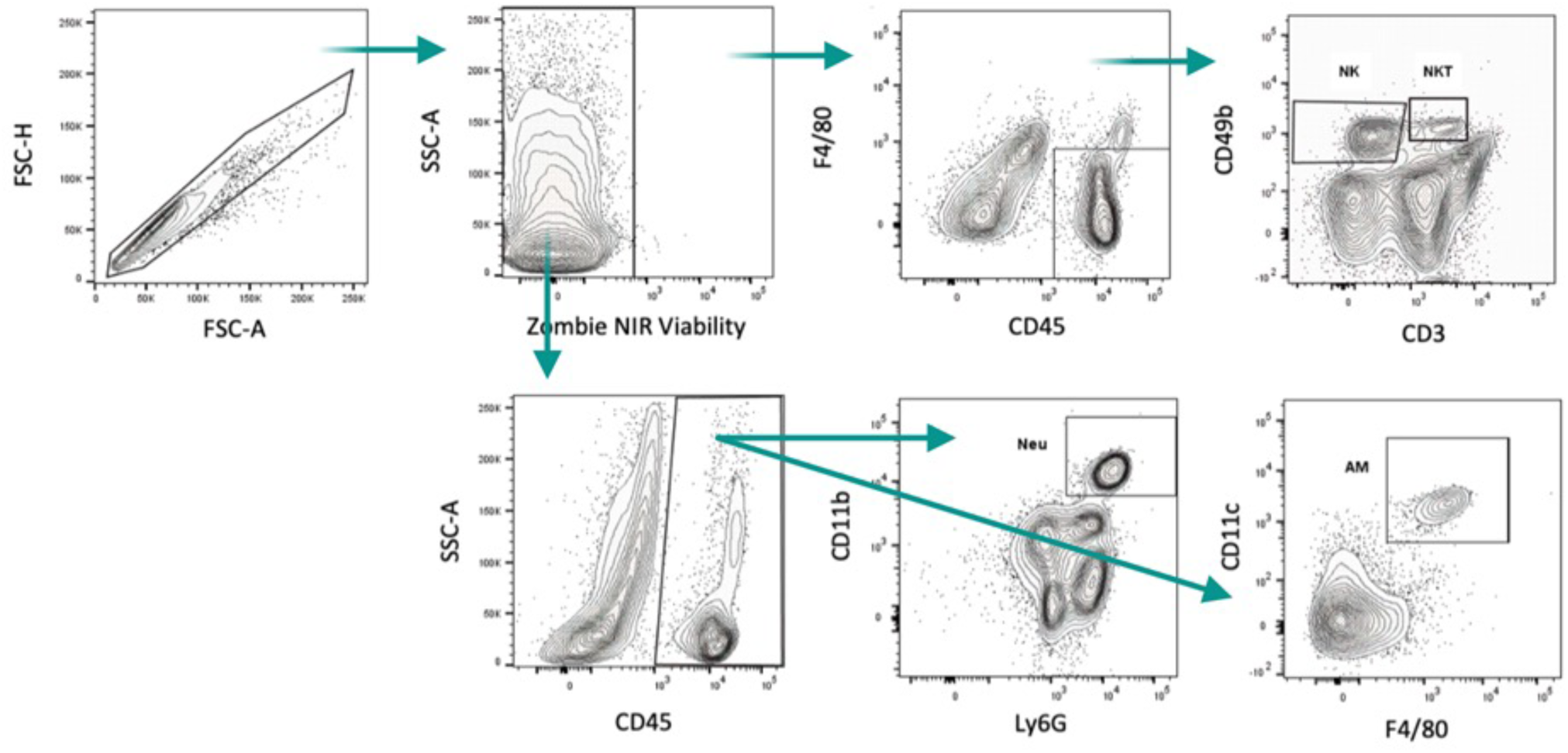
Schematic for analysis of neutrophils, alveolar macrophages (AM), natural killer (NK), and NKT cells by flow cytometry. At days 1 and 2 p.i., whole lung tissue was collected from the respective groups. Single cell suspension was prepared, stained with live/dead cell dye and fluorochrome-conjugated antibodies, followed by flow cytometric analysis. After gating for Live^+^/Dead^−^ followed by F4/80^-^CD45^+^ leukocytes, NK cells were gated on CD3^−^CD49^+^, and NKT cells were gated on CD3^+^CD49^+^. After gating for Live^+^/Dead^−^ followed by CD45^+^ leukocytes, neutrophils were gated on CD11b^+^Ly6G^+^ and AMs were gated on CD11c^+^F4/80^+^.

**Table S1.** Percentages of CD4^+^ and CD8^+^ T cells subpopulations following HIF-1α inhibition at days 4 and 7 post RSV infection. Data from two independent experiments (n = 9-10 mice/group).

|  | RSV/PBS |  |  |  |  |  | RSV/PX478 QAD |  |  |  |  |  |
| --- | --- | --- | --- | --- | --- | --- | --- | --- | --- | --- | --- | --- |
|  | Percentage from CD4<sup>+</sup> |  |  | Percentage from CD8<sup>+</sup> |  |  | Percentage from CD4<sup>+</sup> |  |  | Percentage from CD8<sup>+</sup> |  |  |
|  | Active | Eff | Naive | Active | Eff | Naive | Active | Eff | Naive | Active | Eff | Naive |
| **D4** | 47.3 | 34.9 | 38.1 | 30.4 | 14.1 | 49.2 | 29.4 | 27.3 | 49.5 | 17.1 | 8.1 | 59.9 |
| **D7** | 16.8 | 37.7 | 22.5 | 58.8 | 65.2 | 10.0 | 18.7 | 37.1 | 20.9 | 62.9 | 61.0 | 10.6 |

**Table S2.** Percentages of CD4^+^ and CD8^+^ T cells subpopulations following HIF-2α inhibition at days 4 and 7 post RSV infection. Data from two to three independent experiments (n = 10-14 mice/group).

|  | RSV/CO |  |  |  |  |  | RSV/PT2385 |  |  |  |  |  |
| --- | --- | --- | --- | --- | --- | --- | --- | --- | --- | --- | --- | --- |
|  | Percentage from CD4<sup>+</sup> |  |  | Percentage from CD8<sup>+</sup> |  |  | Percentage from CD4<sup>+</sup> |  |  | Percentage from CD8<sup>+</sup> |  |  |
|  | Active | Eff | Naive | Active | Eff | Naive | Active | Eff | Naive | Active | Eff | Naive |
| **D4** | 11.0 | 23.1 | 45.8 | 17.8 | 17.8 | 45.1 | 10.4 | 24.4 | 46.0 | 17.7 | 19.4 | 43.9 |
| **D7** | 19.3 | 45.6 | 22.3 | 47.6 | 54.4 | 14.0 | 19.0 | 44.8 | 20.5 | 49.7 | 53.4 | 12.3 |

